# A biophysically-detailed model of inter-areal interactions in cortical sensory processing

**DOI:** 10.1101/2024.10.13.618022

**Authors:** Sirio Bolaños-Puchet, Michael W. Reimann

## Abstract

Mechanisms of top-down modulation in sensory perception and their relation to underlying connectivity are not completely understood. We present here a biophysically-detailed computational model of two interconnected cortical areas, representing the first steps in a cortical processing hierarchy, as a tool for potential discovery. The model integrates a large body of data from rodent primary somatosensory cortex and reproduces biological features across multiple scales: from a handful of ion channels defining a diversity of electrical types in hundreds of thousands of morphologically detailed neurons, to local and long-range networks mediated by hundreds of millions of synapses. Notably, long-range connectivity in the model incorporates target lamination patterns associated with feed-forward and feedback pathways. We use the model to study the impact of inter-areal interactions on sensory processing. First, we exhibit a cortico-cortical loop between the two model areas (X and Y), wherein sensory input to area X produces a response with two components in time, the first driven by the stimulus and the second by feedback from area Y. We perform a structural and functional characterization of this loop, finding a differential impact of layer-specific pathways in the feed-forward and feedback directions. Second, we explore stimulus discrimination by presenting four different spatially-segregate stimulus patterns. We observe well-defined temporal sequences of functional cell assembly activation, with stimulus specificity in early but not late assemblies in area X, i.e., in the stimulus-driven component of the response but not in the feedback-driven component. We also find the earliest assembly in area Y to be specific to pairs of patterns, consistent with the topography of connections. Finally, we examine the integration of bottom-up and top-down signals. When presenting a second stimulus coincident with the feedback-driven component, we observe an approximate linear superposition of responses. The implied lack of interaction is consistent with the stochastic and hence naive connectivity in the model, but provides a useful foundation for plasticity mechanisms to learn top-down influences. This work represents a first step in the study of inter-areal interactions with biophysically-detailed simulations.

## 1 Introduction

Connections between areas of the neocortex exhibit regular patterns in their laminar origin and terminations, e.g., in macaque (Rockland & Pandya, 1979), rat (Coogan & Burkhalter, 1990) and mouse (Harris et al., 2019)). This anatomical regularity provides a basis for establishing cortical hierarchies, as a way to explain and predict information flow across cortical areas (Felleman & Van Essen, 1991). In the canonical view, lower-order areas send feed-forward (FF) projections primarily to layer 4 in higher-order areas, and these, in turn, send feedback (FB) projections targeting layers other than layer 4, and most prominently layer 1 in lower-order areas.

A further refinement of this view is the dual counterstream architecture, where two streams of FF / FB exist in supra- and infragranular layers with different distance dependencies (Markov et al., 2014): in supragranular layers, a long-distance feed-forward pathway originating in layer 3 and targeting layer 4 of higher-order areas, and a short-distance feedback pathway originating in L2/3; in infragranular layers, feedback pathways originating from layer 6, targeting layer 6 over short to medium distances and layer 1 over long distances, and a short-distance feed-forward pathway originating in layer 5.

Complementary to this structural perspective, there is a notion of functional relevance of cortical hierarchies, with distinct roles for bottom-up information of sensory origin and top-down information of internal origin. In general, top-down influences imply the impact of complex information represented at higher stages of processing on simpler processes at lower stages, and include sensory context, that would be provided by higher order sensory areas, and behavioral context, that encompasses attention, expectation, perceptual task, and hypothesis testing (Gilbert & Sigman, 2007).

Currently there are two main theories of how the structural and functional perspectives come together in sensory processing (discussed in Schuman et al., 2021; Vezoli et al., 2021. In the representational framework, bottom-up information progresses via feed-forward connections to higher-order areas, where increasingly more complex representations of the input are constructed, and feedback only has a modulatory influence on the activity of lower-order areas. On the other hand, theories of predictive coding posit that top-down signals, representing an internal model or prediction of the sensory input, propagate through feedback connections to lower-order areas, where a comparison is made with their bottom-up inputs and a prediction error is computed; this prediction error is then propagated through feed-forward connections to higher-order areas, and used to update the internal model.

Notwithstanding the advances on this topic in the past few decades, the understanding of feed-forward/feedback connectivity and cortical hierarchy, their fine structural substrate and their functional implications, is still limited. In this work, we introduce a biophysically-detailed model of inter-areal interactions with the aim of better understanding the biophysical substrate of top-down signals and their impact on bottom-up sensory responses.

The model is based on a previous data-driven model of the whole non-barrel rat somatosensory cortex, from which it inherits most structural (Reimann et al., 2022) and functional (Isbister et al., 2023) aspects. The present model is unique, however, in that it represents a system of two cortical areas that interact solely through long-range connections. These model areas represent the first two stages in a hierarchy of cortical sensory processing, with only one of them receiving direct thalamic input, and the other receiving feed-forward input and sending feedback output in turn.

While previous models of interacting cortical areas exist, e.g., the hypothesis-driven population firing rate model of Mejias et al., 2016 and the data-driven point neuron model of Schmidt et al., 2018, this is, to the best of our knowledge, the first large-scale, biophysically-detailed model of inter-areal interactions. Unlike these previous models, and given its computational demands, our model is limited to only two cortical areas, but it enables an initial exploration of the cortico-cortical loop (one step feed-forward, one step feedback) across different levels of description, from single cells to local and long-range circuits.

The manuscript is structured as follows. First, we characterize the structural substrate of a cortico-cortical loop in our model. Second, we exhibit the functional expression of the cortico-cortical loop. Third, we investigate structure-function relations while presenting different stimuli. Finally, we examine the interaction between top-down and bottom-up signals.

## 2 Methods

### 2.1 Virtual connectivity tracing

In the model we have access to the full connectivity matrix, including the number of synapses in a connection. Using this information, together with the locations of cell somas and of individual synapses, it becomes possible to perform a virtual “anterograde tracing” by querying all efferent cells from a given source group, or a virtual “retrograde tracing” by querying all afferent cells on a given target group. That is, we can obtain the spatial location of all cells or synapses participating in the connections between any two groups of cells in the model, being through local or long-range connectivity (separately). If we use a flat view of cell positions (Bolaños-Puchet et al., 2024) and appropriate coloring, we can obtain images of virtual tracings with different “markers” (pixel colors) showing “co-expression” for co-located connections or “sequential expression” across multiple single-synapse steps.

### 2.2 Definition of long-range connectivity

Long-range connectivity in the model was extracted from the base model, and has been previously described (Reimann et al., 2022). However, as this is a main point for the present model, we explain here how it was obtained.

We used a previously published algorithm (Reimann et al., 2019) that generates a micro-connectome constrained by meso-scale data on voxel-wise projection strengths (Knox et al., 2018), and region-to-region connection matrices and laminar termination profiles (Harris et al., 2018). In summary, the data was processed to extract the following: mean synapse densities in target regions; most probable laminar termination profiles for triples of projection type, source and target region; topographical mapping between regions using barycentric coordinates adapted to each region in a flatmap; and projection types of single source neurons. The topographical mapping is fixed and does not change with the direction of the projections. All of these parameters are then expressed in a *recipe file* (Project, 2023) describing each pathway between source and target populations. Source populations consist only of excitatory neurons in a particular layer of a region and may also be split by projection class (e.g. layer 5 intra-telencephalic vs. pyramidal tract). Target populations include all layers in a region and targeting specificity is imposed by the laminar termination profile.

This recipe is then fed to a software algorithm that builds a stochastic micro-connectome instance consistent with the input data. The micro-connectome is described at the level of single source cells making synapses onto spatially constrained groups of neuronal compartments in the target regions, which requires having regions in a 3D atlas filled with neuron morphology reconstructions (multi-compartmental neurons).

The data and algorithm were initially implemented for the mouse isocortex using the Allen Mouse Common Coordinate Framework and its associated flatmap (Wang et al., 2020), and required certain adaptations to be used with the rat model. We generated a flatmap of the rat somatosensory cortex (Bolaños-Puchet et al., 2024) to be used with this algorithm. In terms of input data, given the lack of a similar resource for the rat and the closeness between both rodent species, we used the mouse data unchanged. In fact, we used as starting point the same recipe file (Reimann et al., 2019), and we mapped somatosensory regions from mouse to rat based on corresponding body parts (e.g., S1HL hind-limb region in rat to SSp-ll lower-limb region in mouse, etc.). Then, we modified the coordinates of the triangles defining the barycentric coordinate systems in each region and the spread of the target spatial sampling, as these are related to the scale of the flatmap used. In this way, we obtained a recipe adapted to the rat model, in which we further adjusted the synapse densities to remove those expected from local connections (already accounted for in the separate local connectome). Finally, we ran the algorithm to obtain a long-range micro-connectome of rat non-barrel somatosensory regions (Reimann et al., 2022). By restricting this connectome to the extracted subvolumes used as cortical areas in the present model, we obtain the long-range connectivity between them.

Synaptic delays are computed using a conduction speed of 0.3 m/s for unmyelinated portions of axons (intra-region, Stuart et al., 1997), and a different conduction speed of 3.5 m/s for myelinated portions of axons (inter-region, Simmons and Pearlman, 1983). Unmyelinated portions are approximated by straight lines from the soma of the source neuron to the bottom of L6, and from the location of the synapse to the bottom of L6. The myelinated portion is approximated by a straight line from the location of the source neuron to the location of the synapse.

### 2.3 Definition of model cortical areas

Cortical areas in the model were defined by extracting specific subvolumes from the base model of rat non-barrel somatosensory cortex (Reimann et al., 2022). The size of the subvolumes was similar to that of the released model (Isbister et al., 2024), with seven columns of ∼210 *µ*m radius, and was chosen to have a well-connected central column with surrounding columns to dampen edge artifacts. The location of the subvolumes was chosen based on their inter-connectivity.

We analyzed all pairs of hexagons (single columns) in a hexagonal decomposition of the full model in search for two that were strongly interconnected, had laminar termination profiles that were similar to canonical feed-forward (granular targeting) and feedback connectivity (supra- and infra-granular targeting), and were distant enough that an independent 6-column surround could be extracted for each of them. After two such candidate columns were found, we considered the VPM fiber more strongly innervating the candidate area X, and determined the centroid of all cells targeted by it, which then became the center for defining area X. Then, we performed a virtual anterograde tracing (see Methods 2.1) from the same VPM fiber and determined all cells connected to it one step through long-range projections. We determined the centroid of all such cells, which then became the center for defining area Y. For each of these center positions, a new hexagonal grid was laid out, and the central 7 hexagons in the corresponding volume decomposition were taken as subvolumes representing cortical areas in the model. We then extracted these subvolumes keeping all their local connections, but removing all connections between them. This way, the areas are only connected through long-range connections.

In summary, we performed a selection based on the structure of input connectivity and of connectivity between the areas. As a last step, we verified via virtual anterograde tracing that projection of cells in area Y covered well the central column of area X, thus establishing the structural basis for the cortico-cortical loop. The resulting subvolumes for areas X and Y are contained in regions S1HL (primary somatosensory, hindlimb) and S1FL (primary somatosensory, forelimb), respectively, of the base model. These correspond to mouse regions SSp-ll (primary somatosensory, lower limb) and SSp-ul (primary somatosensory, upper limb), respectively, used in the definition of the long-range connectivity (Reimann et al., 2022).

### 2.4 Spatio-temporal structure of stimuli

Spatial structures of thalamic fibers used as stimulus, all in the central column of area X: random sampling, round spot in the middle, four quadrants, twin spots across halves.

Spike times of thalamic fibers are based on a log-normal fit to an experimentally measured distribution of VPM cell firing (Diamond et al., 1992). Each fiber is assigned a single independent sample from the fitted log-normal distribution. We chose this kind of stimuli over the previously used longer-lasting stimuli with a continuous rate function and involving POm firing (Ecker, Egas Santander, et al., 2024), as we found the latter to produce less sharp responses with longer latencies.

When stimuli are presented multiple times, we use a different random seed in each repetition for the spike time sampling, but the spatial structure remains the same.

### 2.5 Calibration of background activity level and stimulus strength

In order to constrain the functional range of the model to a regime comparable to in vivo data, we performed a calibration of three parameters in order to reproduce observed latencies in L5 (Manita et al., 2015) of the first and second components of evoked responses in area X (∼ 23 ms and ∼ 110 ms, respectively) and the first component of the response in area Y (∼ 80 ms). These parameters are (described in detail in Isbister et al., 2023): *R_OU_*, the ratio of standard deviation to mean of the Ornstein-Uhlenbeck (OU) noise process injected to each cell, *P_F R_*, the fraction of the target per-layer *in vivo* firing rates, and *F_p_*, the fraction of thalamic fibers activated in a stimulus (a measure of stimulus strength).

We ran a series of simulations doing a wide grid search of these parameters using random VPM fibers in the central column of area X. After a workable range was determined with *R_OU_* = 0.3, we performed a refined calibration using a spatially correlated set of fibers (central spot) as stimulus, for all combinations of *F_p_* = 0.15, 0.20, 0.25, 0.30 and *P_F R_* = 0.2, 0.4, 0.5, 0.6, 0.7, 0.8. For each simulation, we computed the mean PSTH of L5 excitatory cells (average of 10 stimulus presentations) from -50 ms to 250 ms after stimulus onset, and performed semi-manual peak detection using the Interactive Peak Finder / Fitter (Octave versions, O’Haver, 2024). We extracted peak amplitudes and latencies, which allowed us to find the parameter combination that best approximated all three latencies of L5 peak firing. The resulting parameter combination is (0.3, 0.6, 0.25).

### 2.6 Circuit manipulations

We performed various circuit manipulations to test the impact of specific pathways present in the connectivity. To this end, we used the capabilities of the Neurodamus simulation control application (Pereira et al., 2024) to specify connection weights between specific groups of neurons. Complete block of a pathway is thus specified by a zero connection weight between select (e.g., layer-wise) neuron populations in areas X and Y.

Additionally, TTX inactivation of sodium channels is modeled using a variable in the sodium channel MOD file that inactivates it completely. This can be easily specified as well for a given population using a Modification section in the simulation configuration file.

In the case where activation of a specific population is desired, mimicking optogenetic rescue of a blocked pathway, we perform current injection in all (or a user-defined fraction of) neurons in the population. Since the threshold current for each neuron is known beforehand, we can inject a step current at a given percent of threshold at the soma of all neurons to make them fire. Other parameters of the stimulation include the start time and duration of the current injection.

### 2.7 Quantification of spiking responses

We use a peri-stimulus time histogram (PSTH) to quantify the level and time-course of evoked activity. We compute them by binning time regularly, counting the number of spikes from a given population in each time bin, and normalizing by the product of the time bin length and the number of cells in the population. This way we obtain mean firing rates as a function of time (up to the resolution of the time bin) for the given population. When a stimulus is presented multiple times under the same conditions (e.g., spontaneous activity level), we perform PSTH averaging to quantify the mean response to the stimulus under those conditions.

Time from stimulus onset refers here to stimulus presentation, since thalamic onset is captured by the firing time distribution of thalamic fibers (see Methods 2.4) and cortical onset can be determined from the peak response of neurons in the model.

### 2.8 Detection of functional cell assemblies

To detect functional cell assemblies of excitatory neurons, we used previously published methods (Ecker, Egas Santander, et al., 2024). In short, significant time bins (with a firing rate above threshold) are hierarchically clustered based on the cosine similarity of their associated spiking vectors (a vector of dimension equal to the total number of spiking neurons and having as entries the number of spikes from each neuron in that time bin). To each cluster of time bins corresponds an assembly, and neurons belonging to an assembly are those whose correlation with the firing of the cluster is above chance level. This means only one assembly is considered active in each time bin, but neurons can belong to more than one assembly.

The cutting threshold for the clustering tree was chosen based on the Davies-Bouldin index, which evaluates to a lower value (better) for higher inter-cluster separation and lower intra-cluster variation. It is also possible, however, to manually specify the number of clusters (and hence the cutting threshold) when this would lead to extracting meaningful information.

We restricted assembly detection to the central column of each area, since these parts of the model are considered well-connected and free from edge artifacts.

### 2.9 Dimensionality reduction

To obtain the spatial structure of each PCA component vector, we computed a histogram of the absolute values of the vector’s components and applied Otsu’s algorithm to threshold the histogram (maximizing inter-cluster variance and minimizing intra-cluster variance). We then applied the same threshold value independently to positive and negative vector components, thus obtaining the identities and locations of significant neurons in the positive and negative parts of each vector. Since the values of the vector components represent firing rates, a positive direction means an increase in firing rate, and a negative direction a decrease in firing rate (relative to mean, in a normalized spiking matrix).

We used the CEBRA algorithm to generate an embedding space to visualize neuronal trajectories after presentation of single or coupled stimuli. The input data to the algorithm was the normalized spike matrix with rows representing time bins (5 ms), and columns (mean-subtracted, unit norm) representing firing neurons. Empty time bins were filtered out. We used the following parameters to train the CEBRA model in CEBRA-time mode:

CEBRA(batch_size=1024, conditional=’time’, device=’cpu’, learning_rate=0.001, max_iterations=20000, min_temperature=0.3, model_architecture=’offset1-model-mse’, num_hidden_units=64, output_dimension=3, temperature=1, temperature_mode=’auto’, time_offsets=5, verbose=True)

We computed mean trajectories (over 40 presentations) for each pattern in the embedding space, and visualized those.

## 3 Results

### 3.1 Structural basis of the cortico-cortical loop

We constructed a biophysically-detailed model of two interacting cortical areas. The two model areas, that we call area X and area Y, represent the first two steps in a hierarchy of cortical sensory processing (Fig. 1A). Area X represents a primary sensory area, as it receives direct thalamocortical (TC) input, whereas area Y represents a higher order sensory area, receiving feed-forward (FF) inputs from area X only. Area Y, in turn, sends feedback (FB) projections back to area X, thus closing the cortico-cortical loop. Additionally, each area receives extrinsic background input representing activity from the rest of the cortex (in general, from the rest of the brain) that is not included in the model.

**Figure 1:**
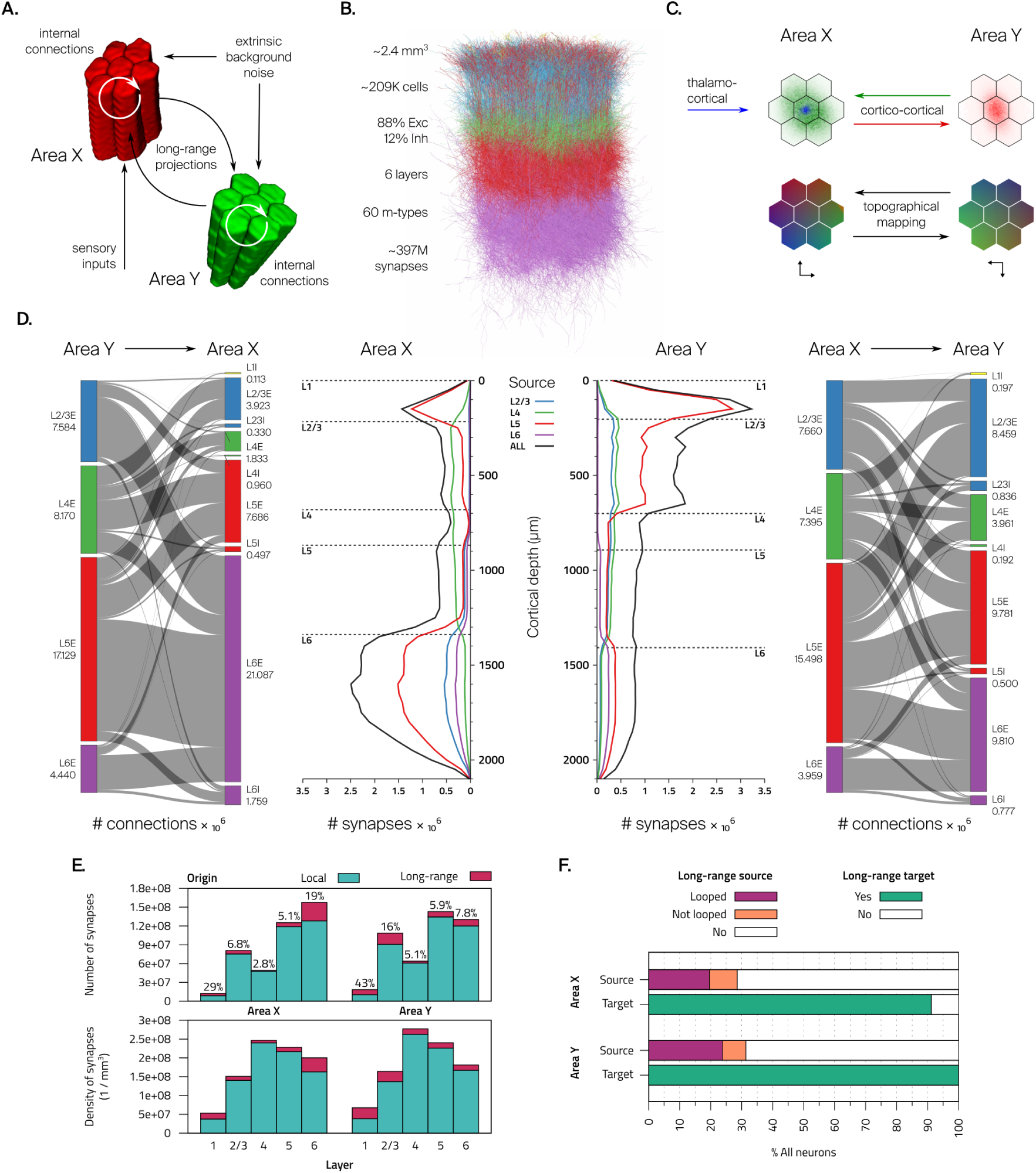
Structural basis of the cortico-cortical loop. **A.** Diagram of connections and input sources in the model. **B.** Three-dimensional rendering of one of the model cortical areas with neuronal morphologies colored by layer (1% of neurons shown, axons not shown). **C.** Top. Virtual anterograde tracing showing reciprocal connectivity. A central TC fiber in area X contacts cells marked in blue, which then project through long-range connections to the cells marked in red in area Y. These cells in turn map back to cells marked in green in area X. Bottom. Topographic mapping between areas, with similar colors mapping to each other. Axes below indicate approximate 180 degree rotation. **D.** Laminar structure of long-range connectivity between model areas. Left. Ribbon map of connections between source and target layer-wise populations and depth-wise histogram of synapses in area X. Right. Same, but for area Y. Color indicates layer. **E.** Top: total number of synapses per layer in each area, from local and long-range origins. Percentages correspond to long-range origin. Bottom: same data, but expressed as synapse densities per layer. **F.** Fraction of neurons that are sources of long-range connectivity (looped or not) and fraction of neurons that are targets of long-range connectivity (at least one synapse), in each area.

Structurally, each area consists of a piece of cortex the size of seven (a central plus six surrounding ones) hexagonal micro-columns with a radius of ∼210 *µ*m each. Area X has 203, 339 neurons in 2.3 mm^3^, and area Y has 214, 696 neurons in 2.48 mm^3^. These volumes were extracted (see Methods 2.3) from a previous model of rat non-barrel somatosensory cortex (the *base model* hereafter, Reimann et al., 2022), and thus have the same composition as the base model. In short, areas are populated with multi-compartmental neurons of 60 different morphological types distributed across 6 layers (Fig. 1B), including thick- and slender-tufted pyramidal cells and several classes of interneurons. Area X is innervated by 4,083 thalamocortical fibers having a characteristic “core”-type (e.g., VPM) laminar termination profile (Meyer et al., 2010).

Long-range connectivity between the model areas is strong and reciprocal, as can be ascertained from a virtual anterograde tracing represented in flat space (Fig. 1C, see Methods 2.1). The tracing shows neurons targeted by a central TC fiber in area X projecting to the center of area Y, and the contacted neurons projecting back to the center of area X. The mapping between the areas is also topographic (Fig. 1C), and exhibits an approximate 180 degree rotation.

In terms of laminar structure (Fig. 1D), area Y receives feed-forward connections to all layers, consistent with observations in rat, (Coogan & Burkhalter, 1990), with synapse counts peaking in layers 1 and 2/3. Synapse numbers of feedback connections in area X peak in layers 1 and 6, with relatively much reduced connections to layers 2/3 and 4, consistent with a canonical feedback connectivity pattern. The pathway structure, however, is more complex than a clear-cut canonical feed-forward vs. feedback system (Felleman & Van Essen, 1991), as it derives from a data-driven long-range connectome based on mouse connectivity data from bulk injections of layer-specific genetically-encoded tracers (Harris et al., 2019, see Methods 2.2). Total numbers of synapses are ∼379 million in area X and ∼416 million in area Y.

A more detailed characterization of long-range connectivity is possible in the model which cannot yet be achieved experimentally, but has been deemed desirable (Douglas & Martin, 2004; Rockland, 2022). In this regard, we measure the fraction of synapses in each layer from local and long-range origins (Fig. 1E). The largest fractions of long-range synapses can be found in layers 1 and 6 in area X (29% and 19%, respectively), and in layers 1 and 2/3 in area Y (43% and 16%, respectively). Other layers in both areas have less than 10% synapses of long-range origin.

We also measure the overlap between long-range projection source and target populations (Fig. 1F). In area X, 32.2% of excitatory neurons (only excitatory neurons are sources of long-range projections in the model, see Methods 2.2) send long-range projections and 91.2% of all neurons receive at least one long-range synapse. In area Y, the corresponding values are 35.5% and 100%, respectively. There are 304, 337 direct loops between projection neurons in both areas (neuron A connects to neuron B, which then connects to neuron A), with 22.1% of excitatory neurons in area X and 27% of excitatory neurons in area Y participating in at least one of them.

### 3.2 Functional expression of the cortico-cortical loop

Physiological details of the model areas are, once more, the same as in the base model (Isbister et al., 2023). In short, neurons are populated with ion channels of Hodgkin-Huxley type for various ionic currents, and the spatial distribution of channel densities is optimized to reproduce firing patterns from experimental electrophysiological traces (Reva et al., 2023). In combination with morphological diversity, this leads to a total of 212 morpho-electrical neuron types. In order to establish a regime of spontaneous activity, we inject each neuron with a noisy conductance signal at the soma, with parameters calibrated to reproduce experimentally observed in vivo layer-wise spontaneous firing rates.

After having established the structural basis for the cortico-cortical loop, we proceeded to test it functionally. To this end, we defined a stimulation protocol where a single stimulus was presented at 1 Hz, comparable to whisker stimulation protocols in rodents (Petersen & Diamond, 2000). The stimulus consisted of a single spike in each of the central 25% of thalamic fibers innervating the central column of area X, with log-normal distributed spike times (see Methods 2.4). The strength of the stimulus (*F_p_*, fraction of stimulated fibers), as well as the levels of background activity in both areas (*P_F R_*, fraction of *in vivo* firing rates), were optimized to attain peak response latencies in L5 comparable to experimental measurements (see Methods 2.5 and, Suppl. Fig. 1).

We recorded spike times of all neurons across both areas and constructed mean PSTHs (see Methods 2.7) across layer-wise populations, with particular attention to L5 excitatory neurons (as in Manita et al., 2015, see Suppl. Fig. 2 for responses in all populations and Suppl. Fig. 3 for responses across cortical depth). After stimulus presentation, we observed a response in area X consisting of two components, a first peak at ∼ 23 ms and a second peak with lower amplitude at ∼ 110 ms after stimulus onset (Fig. 2A). The response in area Y consisted of a first peak at ∼ 80 ms and a second lower amplitude and wider peak at ∼ 120 ms after stimulus onset.

**Figure 2:**
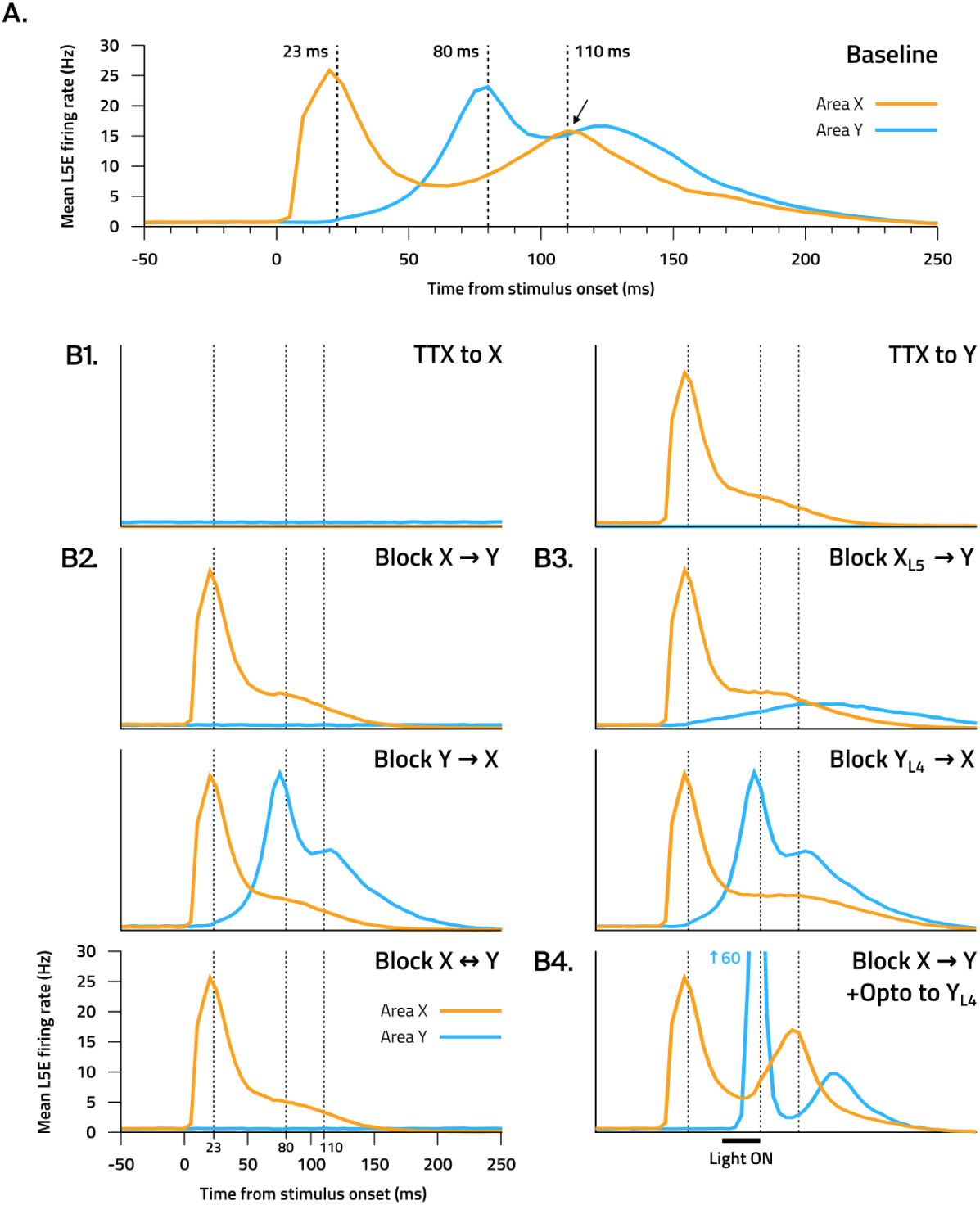
Functional expression of the cortico-cortical loop. **A.** Mean PSTH (N = 30) of L5 excitatory neurons in response to the stimulus in both areas in the baseline condition. It exhibits a second component in the response of area X as a result of feedback from area Y. Dotted lines show experimentally measured latencies (mouse S1 *↔* M2, Manita et al., 2015) to which the stimulus setup was calibrated. **B.** Mean PSTHs (N = 30) of L5 excitatory neurons showing the impact of different circuit manipulations: applying TTX to either area, blocking all connections in either or both directions, blocking L5 in the forward direction and L4 in the backwards direction, and blocking forward connections plus optogenetic stimulus to L4 in area Y. Legend and axes from the bottom left corner plot apply to all plots equally.

In order to characterize the mechanism underlying this response, we performed various virtual manipulations of the system (see Methods 2.6). Initially, we blocked all sodium channels to mimic the action of tetrodotoxin (TTX) application in either of the areas (Fig. 2B1). When applied to area X, the response was abolished entirely in both areas, leaving only the spontaneous firing in area Y. When applied to area Y, the response in area X exhibited only the first component observed before, and was completely silenced in area Y. This pointed out the necessity of area Y firing to obtain a second component in the response of area X.

Then, we performed a series of non-specific blockages of connectivity between areas (Fig. 2B2). Firstly, when blocking all connections from area X to area Y (feed-forward pathway), the response in area X only had the first component observed before, while the response in area Y was completely abolished, leaving only spontaneous firing. This makes sense, since area Y only receives input from area X. Secondly, when blocking all connections from area Y to area X (feedback pathway), we observed a response in area Y consisting of a sharp first peak as observed before, and a second lower amplitude peak, but only the first component in area X. Lastly, when blocking all connections in both directions, the response was the same as when blocking feed-forward connections. These pointed out the necessity of feedback connections from area Y to area X for the appearance of the second component in the response of area X.

Afterwards, we performed blockages of specific neuron populations to study the impact of the layer-wise structure of long-range connections (Fig. 2B3). We show here the pathway blocks that had the greatest effect on the responses (for all pathways, see Suppl. Fig. 4A). Blocking layer 5 connections from area X to area Y (feed-forward pathway), we observe a much diminished response in area Y, and the absence of a second component in the response in area X. On the other hand, blocking layer 4 connections from area Y to area X (feedback pathways) resulted in the suppression of the second peak of the response in area X (except in layers 1 and 6, see Suppl. Fig. 4B). This was surprising, as we’d have expected a stronger impact of blocking L5 than L4, since the Y_L5_ → X pathway is structurally stronger. The impact of blocking a pathway originating from a given layer differed between areas, indicating that in our model these projections have different roles in feed-forward vs feedback pathways, in line with the literature.

Finally, we block all connections from area X to area Y and attempt to rescue the response through optogenetic activation (mimicked as current injection, see Methods 2.6). We observe that providing an appropriately timed optogenetic stimulus of sufficient strength to layer 4 of area Y (90% threshold with 30 ms duration at 50 ms after stimulus onset) produces a high first peak and a second lower amplitude peak in the response of area Y, and a second peak in the response in area X. This points to the sufficiency of activation of layer 4 in area Y (with an intact feedback pathway) to produce a second component in the response of area X.

Altogether, we find that in the model, activation of area Y and the presence of feedback connections are both necessary and sufficient for the existence of a second component in the response in area X. Hereafter, we call this second component the feedback-driven (FD) component, in contrast to the first stimulus-driven (SD) component.

It is notable that the latency of the FD component cannot be explained simply in terms of the synaptic delays of projection neurons across both areas, which average from 3.2 ms for the L6E → L6E pathway to 11.6 ms for the L23E → L23E pathway, in each direction. These delays are considerably shorter than both the ∼ 57 ms between the peak of the SD component in area X and the first peak of the response in area Y (feed-forward latency), and the ∼ 30 ms between the latter and the peak of the FD component in area X (feedback latency), pointing to an important role of local processing and buildup of activity in each area.

### 3.3 Structure-function relations in responses to different stimuli

After examining structural and functional aspects separately, we sought to investigate structure-function relations. To this end, we performed functional assembly detection (see Methods 2.8, Carrillo-Reid et al., 2015; Ecker, Egas Santander, et al., 2024; Herzog et al., 2021) on the evoked activity of excitatory neurons in either area resulting from the presentation of four different stimulus patterns to area X. The stimulus patterns (P_1_ to P_4_) are non-overlapping quadrants that utilize each 25% of the thalamic fibers innervating the central column of area X. They were presented 40 times each in random order at 1 Hz, and produced reliable and well-defined spiking responses in both areas (Fig. 3A).

**Figure 3:**
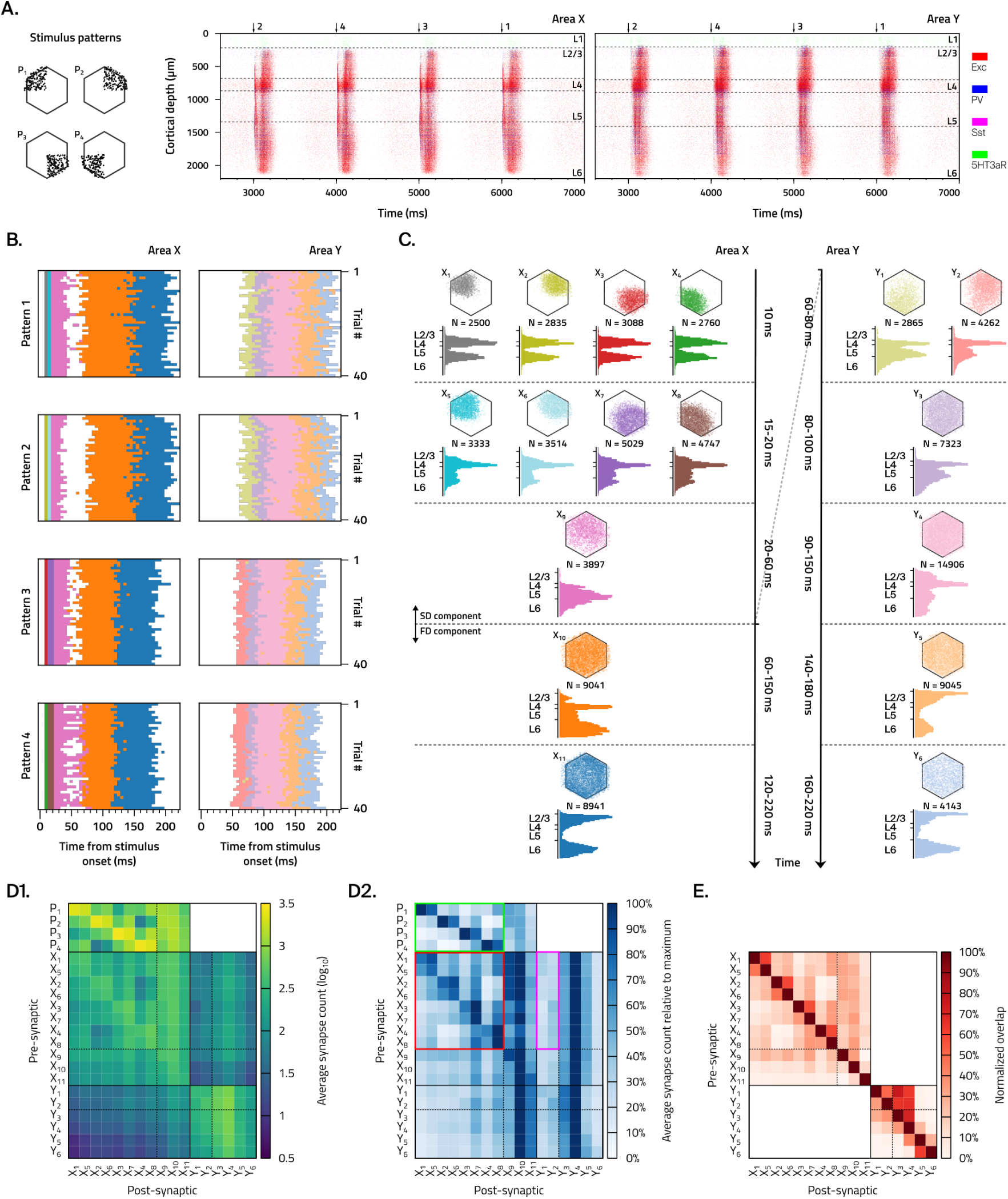
Structure-function relations in responses to different stimuli. **A.** Fibers belonging to each stimulus pattern and raster plots of both areas (left, area X; right, area Y) showing evoked activity from four stimuli (one of each pattern) in the series. **B.** Time course of assembly activation in both areas (left, area X; right, area Y) across all trials (N = 40). Every pixel represents which assembly was active in that time bin (5 ms), with a different color for each assembly. White pixels correspond to no active assembly in those time bins. **C.** Timeline of assembly activation in both areas (left, area X; right, area Y). Each assembly is presented with its color (corresponding to the plots in B), its name (top left), the number of cells belonging to it (middle), and the location of these cells in a flat view (top) and in a depth-wise histogram (bottom). Time flows downwards, and timestamps shown are approximated across all stimulus patterns and trials. The first three rows in area X correspond to the SD component, the last two to the FD component, which is contemporary with the response in area Y. **D1.** Matrix of average synapse counts between assemblies, normalized by the number of pre-synaptic neurons. The first block of rows represents connectivity from the stimulus pattern fibers to the assemblies in area X, with values normalized per fiber. Color scale is logarithmic and assemblies are sorted in temporal order. Thin lines separate blocks of early (stimulus-specific) assemblies in each area. **D2.** Matrix of average synapse counts relative to the maximum value across each row, and separately for each area. Highlighted colored blocks are mentioned in the main text. **E.** Assembly overlap matrix, normalized by size of pre-synaptic assembly.

We found some regularities in the time course of assembly activation for each pattern across all trials (Fig. 3B). First, significant activity (with above-threshold firing rate) lasted no more than 220 ms after stimulus onset in both areas. Second, a similar sequence of active assemblies was seen across trials for each stimulus pattern, but with some pattern-specific differences in early assemblies. Indeed, early activity in area X (first 10-40 ms) consisted in the activation of two short assemblies specific to each stimulus pattern, followed by a longer non-specific assembly. Late activity, present after a short gap (about 60 ms onwards), consisted in the activation of a couple of even longer non-specific assemblies. The early sequence seen here is similar to that previously observed in the absence of feedback connections (Ecker, Egas Santander, et al., 2024), although condensed in time due to our use of shorter-lived stimuli. In that work, early assemblies were pattern-specific, middle assemblies were specific to combinations of patterns (as they were presented), and the late assembly was non-specific. They found no assemblies corresponding to our late assemblies, which confirms their association with feedback inputs. Third, activity in area Y occurred with a delay of about 60 ms, in line with the timing observed in Fig. 2, and showed some variability across patterns. The earliest assembly was pattern-specific (one for P_1_ and P_2_ and another for P_3_ and P_4_), and was followed by a sequence of non-specific late assemblies.

Some insight into structure-function relations can be obtained from a more detailed view of the assembly sequence (Fig. 3C), where we consider the location of the cells belonging to each assembly.

In area X, the first assembly is stimulus pattern-specific (X_1_ for P_1_, X_2_ for P_2_, etc.) and consists of cells whose flat locations largely align with those of the fibers from the corresponding stimulus pattern. The depth location of these cells, peaking in L4 and lower L5, is also consistent with the synapse profile of thalamic innervation. After these initial assemblies, we have another set that is still pattern-specific (X_5_ follows X_1_, X_6_ follows X_2_, etc.), but with larger size, wider flat distribution, and a reduced depth-wise lower peak that spreads more towards L6. The assembly that follows (X_9_) is no longer pattern-specific, and has a flat distribution that is no longer restricted to a quadrant but covers the whole column, with a depth-wise distribution shifted towards infragranular layers. These first three assemblies correspond to the SD component of the response in area X and, as mentioned above, correspond to what has been previously observed in the absence of feedback connections (Ecker, Egas Santander, et al., 2024).

The FD component of the response in area X is described by two assemblies. First is X_10_, a large and long-lasting assembly that covers the whole column, driven by feedback signals from area Y as exhibited by the marked depth-wise peaks in L6, and also L4. The last assembly is X_11_, which is also long-lasting and covers the whole column, but has a depth-wise distribution towards the extremes (L2/3 and L6) with a gap in the middle (L4 and L5).

In area Y, the first assembly is specific to pairs of patterns (Y_1_ for P_1_ and P_2_, Y_2_ for P_3_ and P_4_) and displays some influence from the topography of connections. A shown before (small axes in Fig. 1C), the lower left parts of area X connect to the upper right parts of area Y, which is consistent with the fact that P_1_ and P_2_, with fibers in the upper part of area X, activate Y_4_ with cells in the bottom part of area Y, and the corresponding situation for P_3_, P_4_ and Y_3_. The depth-wise distribution for the first assembly has peaks in L4 and bottom L5. From the second assembly (Y_3_) onwards, pattern-specificity is lost, and activity spreads to the whole column, reaching maximum spread in the longer-lasting Y_4_ that contains over half of the excitatory neurons in the column across all layers, followed by Y_5_ and then Y_6_ with a marked depth-wise distribution towards supra- and infragranular layers.

In an effort to explain the assembly sequence, we looked at the connectivity between all pairs of assemblies in both areas, and also between the thalamic fibers associated with stimulus patterns and assemblies in area X (Fig. 3D). We used as measure of connectivity the average number of synapses between pre- and post-synaptic populations of neurons, normalized by the size of the pre-synaptic population. This allows us to capture an average efferent description of each activated assembly, which may serve to explain the next active assembly in the sequence.

In general, we find strong connectivity of assemblies with themselves, consistent with the notion of assemblies or neuronal ensembles (Yuste et al., 2024), and stronger connectivity between assemblies within the same area than across areas, in line with Fig. 1E.

Average synapse counts (Fig. 3D1) show that the assemblies activated first are indeed strongly innervated by the thalamic fibers of the corresponding stimulus patterns, with slightly lower innervation of the second set of stimulus-specific assemblies. If we consider connectivity relative to the maximum (green block in Fig. 3D2), we also see that these early assemblies are the ones to which the input fibers in each pattern most strongly connect to.

In area X, we find that the two assemblies in each pair of pattern-specific assemblies (e.g. X_1_ and X_5_, X_2_ and X_6_, etc.) are more strongly connected between them than with assemblies in other pairs (red block in Fig. 3D2). This is consistent with the activation of one assembly and then the other in these pairs. Furthermore, we see strong connectivity of all assemblies in area X with X_10_, which may be confounded by the fact that all these assemblies have a considerable overlap with X_10_ (28% ± 6% mean and std. dev., Fig. 3E). Notwithstanding this, activation of X_10_ only happens after activation of area Y. We do not see clear strong connectivity between X_10_ and X_11_, which means the activation of X_11_ is also mediated by the activation of area Y. Indeed, assemblies in area Y connect more strongly to X_10_ and X_11_ among all assemblies in area X, which is consistent as these two assemblies make up the FD component of the response in area X.

Regarding forward connections between area X and area Y, we observe stronger connectivity from assemblies active during the SD component than during the FD component. In particular, we observe strong connectivity to Y_4_, which may be explained by the fact that this assembly covers more than half of the excitatory neurons in area Y, so it’s no surprise to be connected to it. If we look at relative connectivity (magenta block in Fig. 3D2), it is possible to ascertain a relatively larger connection strength between “top” assemblies (X_1_, X_5_, X_2_ and X_6_) and Y_1_, and between “bottom” assemblies (X_3_, X_7_, X_4_ and X_8_) and Y_2_, consistent with the aforementioned topography of connections. This asymmetry underlies the partial stimulus-specificity of early assemblies in area Y. It is interesting, however, that this topographic “priming” of the response of area Y is brief and occurs before X_9_, which is non-specific, but whose activation acts as a “go” signal to trigger area Y mostly through L5 outputs (see its depth-wise distribution in Fig. 3C) that we know are required for area Y activation (see X_L5_ → Y block in Fig. 2).

Finally, within area Y, early assemblies connect strongly to Y_3_ and Y_4_, which again may be explained by their large size and considerable overlap. This is consistent with the order of activation of these assemblies. The transition from Y_4_ to Y_5_ and then Y_6_, however, does not seem readily explainable based on connectivity alone, even when considering connections from assemblies in both areas.

### 3.4 Bottom-up and top-down interactions

After studying the activity evoked by the presentation of a single stimulus, we addressed the topic of interactions between multiple bottom-up and top-down signals.

Initially, we looked briefly at the interaction between two bottom-up signals. We compared the response to separate weak stimuli A and B (10% fibers each), and to the combined stimulus A + B (Fig. 4A). These stimuli are considered weak because they fail to elicit an FD component in area X by themselves. However, when presented together, the response shows a clear FD component. When increasing the delay between A and B in the combined stimulus, we observed a decrease in the amplitude and an increase in the latency of the FD component, to the point where it disappeared (at a delay of ∼ 110 ms). This shows how the presence of an FD component can be a signature of temporal integration of stimuli within a narrow time window.

**Figure 4:**
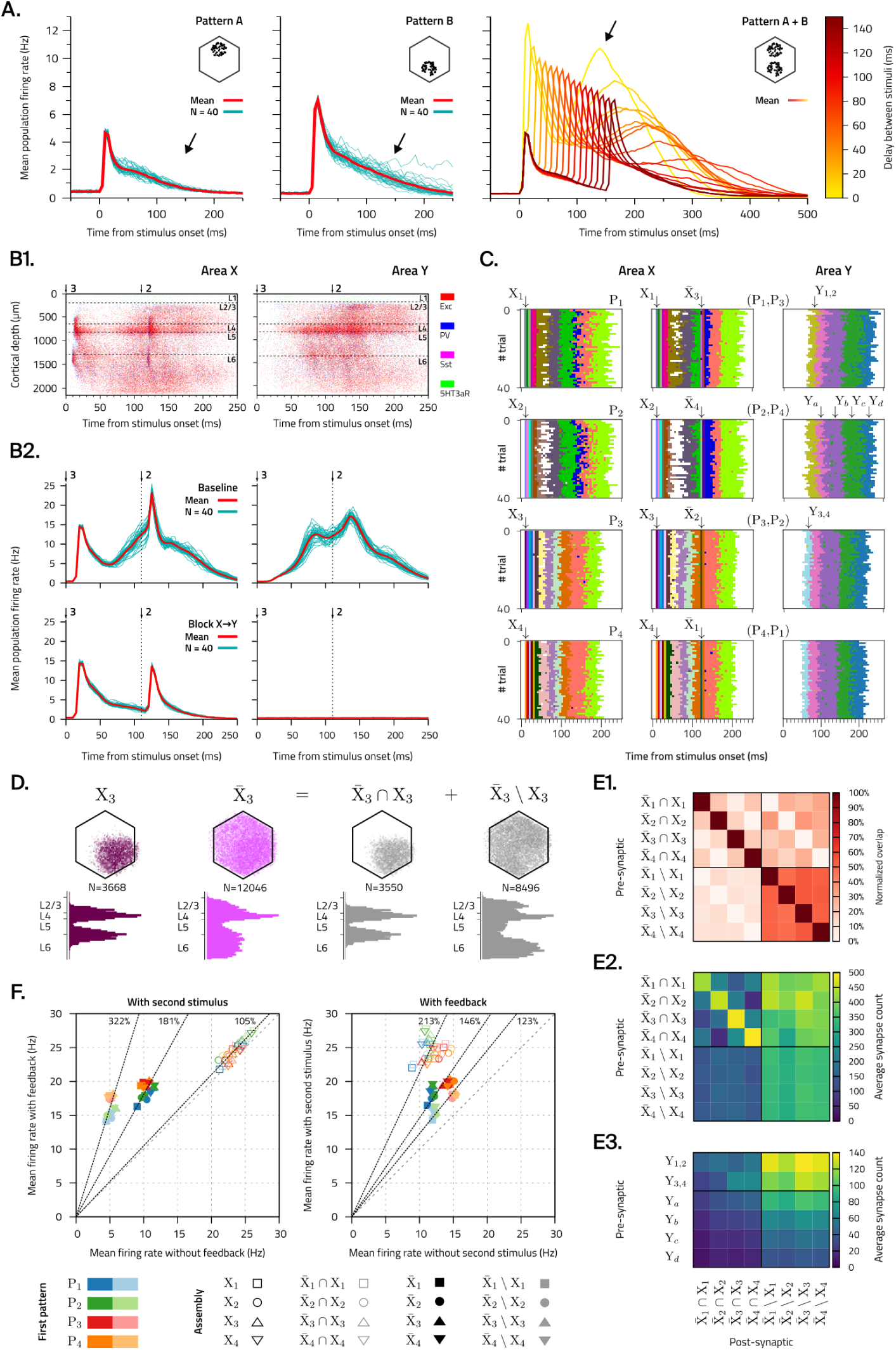
Bottom-up and top-down interactions. **A.** Mean population responses in area X to individual weak stimuli A and B, and to the combined presentation of A + B with a delay in between. We observe an FD component in the response to the combined stimulus (black arrows), but not to the separate stimuli. **B1.** Raster plots of the evoked response in both areas to the presentation of an example stimulus pair (P_3_,P_2_), up to 250 ms after first stimulus onset. **B2.** Mean population PSTHs (N=40, red) in both areas for an example stimulus pair (P_3_,P_2_) in the baseline (top) and blocked (bottom) conditions. Thin lines (turquoise) represent individual trials. **C.** Assembly activation sequences in area X after presentation of individual patterns (left column) or pairs of patterns (right column). Pattern-specific assemblies activated by the second stimuli (X̅ *i*) are different than for the first stimuli (X*_i_*). Colors represent different assemblies (43 in total). **D.** Numbers, flat locations and depth-wise histograms of neurons in the pattern-specific assemblies active after the first (X_3_) and second (X̅ 3) stimulus presentation of the example pattern P_3_, as well as their intersection (X_3_ *∩* X̅ 3) and difference (X_3_ *\* X̅ 3). **E1.** Normalized overlap between assembly subsets active in area X. **E2.** Connectivity matrix of assembly subsets active in area X between themselves, and **E3.** from assemblies active in area Y. Normalized by the number of pre-synaptic neurons. **F.** Change in mean firing rate of neurons in pattern-specific assemblies to the first and second stimulus in the time window from t=110 ms to 140 ms after stimulus onset. Left: With second stimulus, across baseline and blocked conditions. Right: In the baseline condition, with or without a second stimulus. Colors indicate the pattern of the first stimulus, with dark and light shades indicating full or subset assemblies.

A more interesting matter, however, is studying the interaction between the presented stimulus (bottom-up) and the ongoing feedback-derived (top-down) activity. We dedicate the rest of the section to this topic, and try to answer the following question: how is the response to a stimulus different when presented in isolation and when presented during an ongoing feedback-derived response?

To this end, we defined a stimulus protocol using pairs of the same four patterns defined in the previous section. The second stimulus in a pair is presented with a delay of 110 ms, making it coincide with the FD component so the interaction between bottom-up and top-down signals can take place. This delay also ensures that the stimuli are essentially independent, in accordance with the observation above. We tested all sixteen pairs of the four patterns, e.g. (P_1_,P_1_), (P_1_,P_2_), (P_1_,P_3_), etc., and presented each stimulus pair 40 times in random order at 1 Hz. As a control, we ran an equivalent set of simulations with forward connections blocked between area X and area Y (”blocked” condition), as well as additional simulations presenting only the first or only the second stimulus in each pair.

The first thing we observed is that the second stimulus in a pair produces a sharp transient response on top of the FD component of the response to the first stimulus. This is visible in both the spike rasters (Fig. 4B1) and in the mean population PSTHs (Fig. 4B2 top), shown here for an example stimulus pair (P_3_,P_2_). In the PSTHs we notice a new peak in the response of area X that adds to the FD component, with higher amplitude than the peak from the first stimulus. In area Y, we find likewise an increased amplitude in the second peak of the response. It is notable that the presentation of the second stimulus does not trigger a subsequent FD component of its own, i.e., a third component in the response, but is rather “absorbed” by the ongoing response to the first stimulus. In the simulations with blocked forward connections between area X and Y (4B2 bottom), we observe no evoked activity in area Y, as expected, and thus no FD component in the response of area X to the first stimulus. This results in both stimuli in a pair producing essentially independent responses.

Next, we looked at assembly activation sequences in area X, using all simulations in both the baseline and blocked conditions for functional assembly detection (in total over 67 million spikes in about 30 minutes of biological time). When a single stimulus is presented (Fig. 4C left column), we observe pattern-specific early assemblies (as in Fig. 3B), and two sets of assemblies during the FD component, earlier ones that have specificity for patterns P_1_ and P_2_ vs P_3_ and P_4_, and later non-specific ones. When presenting a second stimulus (Fig. 4C middle column), we observe brief activation of a pattern-specific assembly in the middle of the FD component (at 120 ms after stimulus onset), and otherwise the assembly sequence is unchanged. These results differ from what we observed before (Fig. 3B) as we chose here a lower clustering threshold than the default value (computed by minimizing the Davies-Bouldin index, see Methods 2.8), in order to have a finer look into the impact of the second stimulus. In area Y (Fig. 4C right column) we keep the default clustering threshold, and observe the same as before: an initial assembly specific to pairs of patterns (Y_12_ and Y_34_) followed by a sequence of non-specific assemblies (Y*_a_* to Y*_d_*).

Indeed, we lowered the clustering threshold (increasing the number of clusters) to the point where the assemblies activated by the first (X_1_ to X_4_) and second stimuli (X̅ _1_ to X̅ _4_) were different. This choice, albeit arbitrary, allowed us to identify relevant groups of neurons that we analyzed further. A comparison between the two sets of pattern-specific assemblies (Fig. 4D, shown for P_3_; for all patterns see Suppl. Fig. 5A) shows that the assemblies active during the second stimulus have about three times more cells, are more spread throughout the column, and have a significant contribution from L6, in comparison to the assemblies activated by an isolated stimulus.

Furthermore, the X̅ *_i_* can be decomposed as sets into their intersection with X*_i_* (X̅ *_i_* ∩ X*_i_* = neurons in both X*_i_* and X̅ *_i_*) and their difference with X*_i_* (X̅ *_i_* \ X*_i_* = neurons in X̅ *_i_* but not in X*_i_*). We observe (Fig. 4D, shown for P_3_; for all patterns see Suppl. Fig. 5B) that each X̅ *_i_* ∩ X*_i_* is essentially the same as X*_i_* (over 96% overlap in all cases), and the X̅ *_i_* \ X*_i_* have a very similar depth-wise distribution and considerable overlap (about 50% in all cases) with the assembly active during the FD component when only a single stimulus is presented (X_10_ in Fig. 3C). Additionally, we observe strong overlap (52% on average, Fig. 4E1) and homogeneous connectivity (bottom right block in Fig. 4E2) of the assembly differences among themselves. Active assemblies in area Y (Fig.4C right column), especially the early assemblies specific to pairs of patterns (Y_12_ and Y_34_), are more strongly connected to neurons in the assembly differences than in the assembly intersections (Fig.4E3), consistent with the feedback-like depth-wise profile.

We further analyze the activity of neurons belonging to these assemblies, by computing their mean firing rate in the time window from 110 to 140 ms after stimulus onset, i.e., after presentation of the second stimulus of a pair, across baseline and blocked conditions (Fig. 4F left). Results across all stimulus pairs, where we analyze the pattern-specific assembly of the second stimulus, show that the mean firing rate of neurons in X assemblies is greater than or equal in the baseline condition than in the blocked condition, but by a small amount (mean 5% higher). This means that neurons active upon stimulus presentation in absence of the FD component do not fire on average considerably more than when the FD component is present. In contrast, the mean firing rate of neurons in X̅ assemblies, is on average 81% higher when the FD component is present than when it is absent. This means that this second group of assemblies better captures the impact of the FD component on the response to the stimulus.

However, when considering assembly subsets, we observe that neurons in assembly intersections do not fire considerably more (5% higher on average) in the baseline than in the blocked condition. On the other hand, neurons in assembly differences fire on average 3.22 times more in the baseline condition than in the blocked condition, meaning these neurons are specifically active during the FD component. Conversely, if we compare the firing rates when the second stimulus is presented to when it is not (Fig. 4F right), neurons in assembly intersections fire on average 2.13 times more, whereas neurons in assembly differences fire comparatively only slightly more (23% higher on average).

Altogether, these results indicate that the response to the second stimulus presented at the same time as the FD component is short-lived (∼ 10*ms*, same as an isolated stimulus) and largely additive. This means that the response is an approximate linear superposition of the activity of neurons (in assembly differences) preferentially responding to the feedback input from area Y and the activity of neurons (in assembly intersections) preferentially responding to the activation of thalamic fibers, with no clear interaction between these two populations. This is consistent with the connectivity between them (Fig. 4E2), where we observe that assembly intersections are well connected to assembly differences (which explains Fig. 4F right), but the reverse is not true, so the activation of neurons in assembly differences has little impact on the response of neurons in assembly intersections (as shown in Fig. 4F left).

In order to gain an understanding of the global response of area X, and to mitigate the risk of becoming biased by looking only at he pactivity of specific sub-populations (the attern-specific assemblies X*_i_* and X̅ *_i_*), we analyzed the activity of all excitatory neurons in the central column of area X using a classical dimensionality reduction technique. As such, we performed principal component analysis (PCA) on the spiking activity of 24,884 excitatory neurons in area X across all simulations with single and paired stimuli in both the baseline and blocked conditions (N=1760 trials in total). We then analyzed global activity trajectories in the first four PCA components (ordered by explained variance).

When single patterns are presented, we observe a clear distinction of responses, with one characteristic direction of deflection for each pattern in the plane of the PCA_1_ and PCA_2_ components (Fig. 5A1 left column in each condition). When stimulus pairs are presented we observe two deflections, each being in the direction corresponding to the pattern presented (Fig. 5A1 right column in each condition). This means stimulus identity is clearly captured by these PCA components and is not lost in the interaction with the FD component. Actually, the pattern-specific directions are preserved in the blocked condition (Fig. 5A1), and when presenting second stimuli interpolated between two patterns we observe intermediate directions (Fig. 5A2), meaning these two PCA components only represent stimulus identity. This is further supported by looking at the spatial structure of the PCA_1_ and PCA_2_ components (Fig. 5C top), which partition the flat view into roughly up/down and left/right parts, and have a depth-wise distribution largely similar to that of thalamic innervation and the initial pattern-specific assemblies.

**Figure 5:**
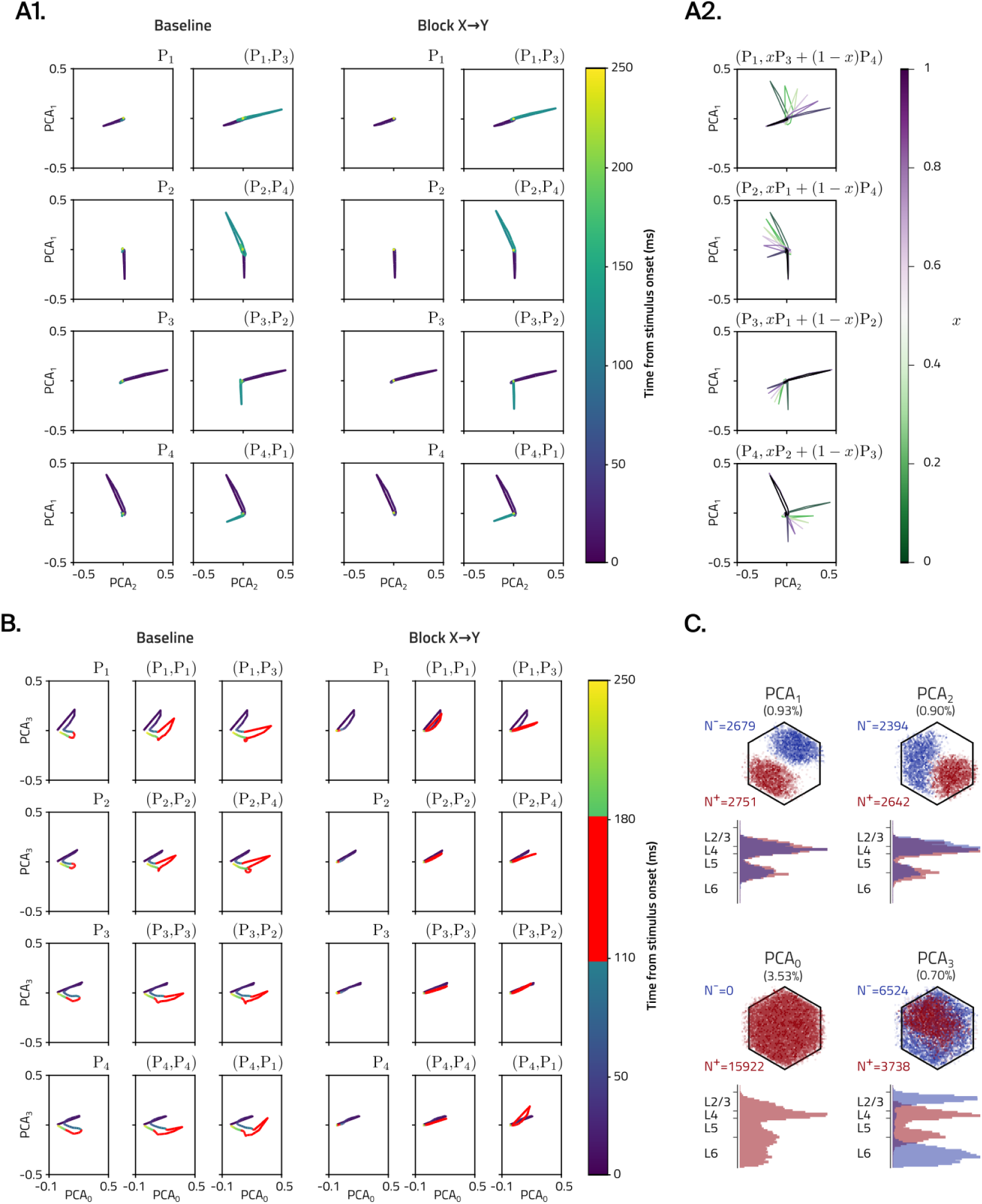
Linear decomposition of global network activity after stimulus presentation. **A1.** Network trajectories in PCA_2_-PCA_1_ space for baseline and blocked conditions after single (left column) and paired (right column) stimulus presentation. The colorbar describes time along the trajectories. **A2.** Network trajectories in PCA_2_-PCA_1_ space for paired stimulus presentations where the second stimulus is a linear combination of two patterns. The colorbar describes the linear mix. **B.** Network trajectories in PCA_0_-PCA_3_ space for baseline and blocked conditions after single (left column) and paired (right column) stimulus presentation. The colorbar describes time along the trajectories, with highlighted time window (red) of feedback interaction. **C.** Locations and depth-wise histograms of significant neurons in the first four PCA vectors with positive (red) and negative (blue) components.

We obtain a richer perspective by further looking at the PCA_0_ and PCA_3_ components. From their spatial structure (Fig. 4C bottom), we can see that PCA_0_ contains a large fraction of the excitatory neurons in the column (64%), with a nonspecific distribution in the flat view and across depth; it also has only positive components, so this PCA component represents more or less global activity. On the other hand, PCA_3_ has a distinct depth-wise distribution that separates “inner” layers (bottom L2/3, L4 and bottom L5) from “outer” layers (top L2/3 and L6), roughly capturing the difference between input and non-input (feedback) participating neurons. This interpretation is consistent with the trajectories of the responses to single or paired stimuli. For single stimuli (Fig. 5B left column in each condition), we observe trajectories with an initial deflection towards positive PCA_3_ and PCA_0_ upon stimulus presentation, followed by a second shorter deflection towards negative PCA_3_ and positive PCA_0_ during the FD component (red highlighted part), before returning to the initial point. For paired stimuli (Fig. 5B middle and right columns in each condition), we observe the same first deflection, followed by a second deflection that moves again towards positive PCA_3_ and PCA_0_ (to higher PCA_0_ than the first deflection, meaning higher firing rate as observed already in Fig. 4B2) on presentation of the second stimulus, and then going towards negative PCA_3_ and PCA_0_ before returning to the initial point. In simulations in the blocked condition (Fig. 5B), the second deflection happens along the same direction as the first (no motion towards negative PCA_3_), as there is no feedback input that would activate neurons in the negative part of the PCA_3_ vector (upper L2/3 and L6).

When showing pairs of the same stimuli (Fig. 5B middle column in each condition) and of different stimuli (Fig. 5B right column in each condition), we find slight differences in the shape of the second deflection, which points to some effect of the identity of the first stimulus (and thus of the FD component) on how the activity unfolds upon presentation of the second stimulus. A related effect was already evident in Fig. 4F, where the spread of points showed some variability in the firing rates for different first stimulus patterns. Furthermore, this is also consistent with the detection of different initial active assemblies in the response of area Y (Fig. 4C right column) for patterns P_1_ and P_2_ (Y_12_) vs P_3_ and P_4_ (Y_34_). The pattern-specific assemblies associated with the second stimulus (X̅ *_i_*), however, did not depend on the identity of the first stimulus (at the default clustering level). Beyond these observations, we do not pursue this matter further.

All in all, the PCA analysis above shows that each stimulus pair generates a network trajectory with clear features in the first four PCA components, that is slightly different for each pair, but that nevertheless follows the same basic plan: starts and ends in the same place and differs only during the time where the FD component is present. In this time window, the interaction between ongoing activity and the second stimulus is seen to be largely additive in both the PCA spaces analyzed above, with the combined trajectories (red parts in Fig. 5B) being to a first approximation the sum of the individual trajectories of the FD component and the second stimulus. It is not evident, thus, how the top-down activity impacts the bottom-up response, but it is clear that the second stimulus does not steer the course of ongoing activity, but briefly adds to it.

## 4 Discussion

We have demonstrated a biophysically-detailed model of inter-areal interactions between two model cortical areas (X and Y). The areas have similar composition, in terms of single-cell models, synaptic physiology, local connectivity rules, etc. and differ mainly in their long-range connectivity. Area X represents a primary sensory area, receiving sensory inputs through thalamic innervation and feedback inputs from area Y, while area Y represents a higher-order area, receiving feed-forward inputs from area X only. The hierarchical relation between the areas is consistent with the position in the cortical hierarchy (Harris et al., 2019) of the mouse regions underlying the connectivity in the base model (SSp-ll vs SSp-ul, see Methods 2.3). Additionally, area Y is slightly larger as a consequence of cortical thickness variations in the atlas used to build the base model. This is unintentional but incidentally consistent with observed differences in cortical thickness between primary visual and higher-order visual areas in humans (Alvarez et al., 2019).

We have exhibited a well-defined cortico-cortical loop between the two areas, wherein thalamic inputs to area X produce an activation of area X that propagates to area Y through feed-forward pathways, and then comes back to area X through feedback pathways. Notably, we have reproduced peak latencies in L5 firing (Manita et al., 2015) and observed responses with comparable amplitude in both areas. This behavior was found to be robust, showing a gradual variation in peak latencies and amplitudes of the responses (see Suppl. Fig. 1) as a function of stimulus strength and level of background activity.

We would like to emphasize how remarkable it is to have observed such a well-defined cortico-cortical loop with realistic peak latencies and similar amplitudes in both areas. That is, considering that the model consists of a statistical instantiation generated in a bottom-up manner from experimental data and algorithms based on experimental data, and its behavior emerges from the interactions between all its components across multiple scales, constrained by data on basic biophysical features (synapse densities, firing patterns, PSP amplitudes, spontaneous firing rates, etc.) without tuning to any particular higher-order function.

We have analyzed some structural and functional aspects of the cortico-cortical loop. In particular, we have found a differential impact of layer-specific pathways, with prominent roles for L5 activation in area X and L4 activation in area Y. We have also observed well-defined temporal sequences of functional cell assembly activation. Upon presentation of four different stimulus patterns, early assemblies of the stimulus-derived (SD) component in area X were found to be pattern-specific (in line with Ecker, Egas Santander, et al., 2024), whereas late assemblies of the SD component and assemblies of the feedback-derived (FD) component were non-specific. In area Y the first assembly was found to be specific to pairs of patterns, thus demonstrating that a stimulus-specific response can be preserved even along a naive long-range pathway that was statistically instantiated, owing to the topography of the underlying connections.

Finally, we have used the model to study the role of inter-areal interactions in cortical sensory processing. We have sought to understand how top-down signals in the FD component interact with bottom-up signals from a second stimulus presented simultaneously. Using techniques of assembly detection and dimensionality reduction, we have found evidence of an approximate linear superposition between top-down and bottom-up activity. This additivity implies a lack of interaction between top-down signals and bottom-up responses to sensory stimuli, consistent with the observed weak coupling between the neuron populations that receive top-down and bottom-up inputs (Fig. 4E2). We think this is a consequence of the connectome being a statistical instantiation, meaning that the network is initially naive, and the lack of plasticity mechanisms in the model (other than short-term synaptic plasticity, M. V. Tsodyks and Markram, 1997), so the network stays naive throughout simulations. The fact that both signal streams are represented without interfering destructively with each other, however, is a conducive foundation for the formation of associations if plasticity was indeed present. Put another way, a stronger interaction between signal streams in the naive network could have severely curtailed the potential for plasticity.

### Comparison to previous models

In comparison with the base model, even though all components were already present and some analyses had been conducted on the structure (Fig. 8 of Reimann et al., 2022) and function (Fig. 8 of Isbister et al., 2023) of long-range connections, the model has exhibited previously unseen behavior thanks to its reduced setting tailored to observe the impact of long-range connectivity. Having two separate areas with no other cortical inputs (other than background noise) and no other connectivity (other than internal), it represents a minimal system to study inter-areal interactions with biophysical detail.

In contrast, previous models with less biophysical detail have addressed other aspects relevant to cortical function. We describe below two models from the literature, and highlight some differences to our model.

The model of Schmidt et al., 2018 consists of 32 interconnected areas involved in visual processing and is based mainly on macaque data. This model uses a microcircuit to model each area, with 8 populations (one excitatory and one inhibitory for each of L2/3, L4, L5 and L6) of leaky integrate-and-fire neurons per area; in comparison, we have 9 populations (also L1) of multi-compartmental neurons with Hodgkin-Huxley type ion channels. Connectivity in their model considers both local and long-range connections and is generated by random sampling of pre- and post-synaptic pairs, using per-area and per-population connection probabilities, with inter-areal layer specificity based on layer-specific tracing data and synapse densities; in our case, we have touch-based local connectivity and data-driven long-range connectivity with specific laminar termination profiles. Beyond these differences in model composition and level of description, the main difference with our work is their focus on spontaneous activity manifested by the network under random noise inputs, mediated by inter-areal connections between all areas. We focus instead on evoked activity resulting from the activation of thalamic fibers in one of two areas, and consider spontaneous activity as part of the base model (Isbister et al., 2023). We only dealt explicitly with spontaneous activity when calibrating the latency of peak responses through the *P_F R_* parameter of the per-layer noisy conductance injection (see Methods 2.5 and Suppl. Fig. 1).

The model of Mejias et al., 2016 similarly consists of 30 interconnected areas involved in the macaque visual hierarchy. In this model each area is represented by 4 populations (one excitatory and one inhibitory for each of supragranular L2/3 and infragranular L5/6) whose mean activity is described by a nonlinear firing rate model of Wilson-Cowan type. Connectivity is described by a connectivity matrix specifying the weights of the coupling between different populations across areas, and is specified at four levels: intra-laminar, inter-laminar, pair-wise inter-areal (laminar termination pattern) and large-scale (hierarchical position). Once more, beyond these differences in model composition and level of description, the main difference with our work is their focus on the oscillatory behavior of neuron populations within an explicit functional hierarchy. In our model, we only have two areas and thus no possibility of establishing a complete hierarchy. Furthermore, the hierarchical relation between the two areas in our model is not imposed but stems purely from the structure of the connectivity, with only one of them receiving sensory inputs, and the connections between them having different laminar profiles. We also did not consider oscillations nor their relation to feed-forward and feedback activity, a topic which has been recently criticized in Vinck et al., 2023. The only frequency-related parameter we considered was the frequency of stimulus presentation, which was kept fixed at 1 Hz. Nonetheless, in supplementary simulations with higher stimulus frequencies (Suppl. Fig. 11), we found that at frequencies higher than 2 Hz the FD component disappeared for all but the first stimulus presented. While the response in area X did follow the frequency of stimulation, its amplitude was reduced the higher the frequency, until it failed to produce a response in area Y, which in turn resulted in the disappearance of the FD component in area X. The first stimulus in the series, however, was always effective. This can be explained by the network being in a fully recovered state at the start of the stimulus series, and in a partially depleted state (a combination of synaptic vesicle depletion and neuron refractoriness) throughout the stimulus series, with less time to recover the higher the frequency.

### Beyond the canonical feed-forward/feedback model

Being built according to a data-driven model (Harris et al., 2018; Knox et al., 2018; Reimann et al., 2019), the connectivity between the two regions showed some discrepancies with respect to the canonical view of feed-forward/feedback connectivity (feed-forward terminates in L4 and feedback terminates elsewhere than layer 4 Felleman and Van Essen, 1991). First, the existence of a peak in L1 in the laminar termination profile for feed-forward projections (see Fig. 1D), and second, the relevance of feedback pathways originating in L4 for the generation of the FD component in area X (see Fig. 2B3).

Regarding the first point, it should be noted that this peak is formed mainly by synapses from L5 pyramidal cells in area X onto the apicals of L5 pyramidal cells in area Y. This is explained because the data-driven long-range connectome specifies a laminar termination profile with increased density towards upper layers (profile 1 of Reimann et al., 2019) and L5 apicals get picked more often by random sampling, since they are much more numerous in L1 than local interneurons. This “superficial-only” pattern was identified as feed-forward in the source publication of long-range connectivity data (Harris et al., 2019).

Regarding the second point, a recent study has found evidence of L4 participation in a non-canonical feedback pathway (Minamisawa et al., 2018), which allows the possibility that our model captures such a relevance of L4 for feedback activity. A similar point was made with regards to structure in the source publication of long-range connectivity data (Harris et al., 2019), where long-range projecting L4 neurons were also directly identified.

It should be noted in this respect that, when we tested alternative connectomes having a more canonical subset of pathways (results not shown), we observed only a very diminished cortico-cortical loop that required higher global excitability conditions ([Ca]*_out_* = 1.1 mM instead of 1.05 mM). All of this is indicative that the complexity of data-driven connectivity goes beyond what the canonical view of feed-forward vs feedback can make sense of, as has been previously noted (Harris et al., 2019). We did find, however, that the effect of blocking layer-specific pathways was different for the feed-forward and feedback directions, in line with the idea that layers play specific roles in hierarchical interactions (Felleman & Van Essen, 1991).

### Analysis considerations

We have sought to understand how bottom-up, stimulus-derived activity interacts with top-down, feedback-derived activity. In particular, we have analyzed neuronal activity in a time window where a stimulus coincided with the FD component from an earlier stimulus, and employed different analysis techniques to determine how the response differs from that of an isolated stimulus (without FD component). We discuss below about the analysis techniques employed.

First, we computed peri-stimulus time histograms (PSTHs) of neuronal firing across layers, depths and areas. These allowed us to find latencies and amplitudes of peak firing (e.g., Suppl. Fig. 1) and to observe the propagation of activity across layers and areas (e.g., Fig. 2 and Suppl. Fig. 2), and along depth (e.g., Suppl. Fig. 3). While PSTHs can provide useful information on the firing of specific populations of neurons, the choice of population has to be made before computing a PSTH, and it is up to us to find functionally relevant populations.

In this regard, functional cell assembly detection provides an automated method to find such functionally relevant populations, in the form of cell assemblies made up of neurons that consistently fire together. This technique allows us to see the time course of activity through the eyes of assembly activation sequences (e.g., Fig. 3B). While these do not provide a quantification of activity, they serve to identify which populations of neurons were significantly active as time goes by. Furthermore, knowing the composition of cell assemblies allows us to establish a structural correlate for the assembly sequence in terms of where the active neurons are (e.g., Fig. 3C, shown here in a flat space + depth representation) and what is the connectivity between them (e.g., Fig. 3D).

One aspect of the assembly detection algorithm that warrants further discussion is the choice of clustering threshold. Even while the clustering threshold can be chosen automatically using some criterion (such as minimizing the Bouldin-Davies index, as is done by default), in the end the choice is, as with every clustering algorithm, arbitrary. This does not mean that it is meaningless, though. Lowering the threshold breaks up clusters, and makes evident finer differences between the assemblies associated with them. Importantly, we do not control how the clusters break up, so it is not the same as picking neurons arbitrarily. In our case, we lowered the threshold to a point where the general structure of the assembly sequence was still preserved across trials (vertical strip structure in Fig. 4C), and only up to the point where the assembly of interest split in two (Fig. 4D). Ultimately, cell assembly detection does not provide a definitive answer, but rather serves to identify relevant neuron populations on which further analyses can be performed (e.g., Fig. 4F).

Finally, we employed dimensionality reduction to analyze the global activity of the whole excitatory population of area X. In contrast to the sub-population view of assemblies, PCA allows us to represent the activity of entire populations of neurons (about 25K in our case) in a comprehensible and interpretable way. In our case, the first four PCA components were very informative of stimulus presentation and network conditions (baseline vs blocked) and provided clearly interpretable trajectories (Fig. 5A and B). All the more after establishing a structural link for the PCA directions (Fig. 5C) by considering the populations of neurons primarily represented by them. We attribute the success of our PCA decomposition to the large amount of ground truth data that we collected over hundreds of simulations (1760 trials and 686 million spikes in total).

Beyond how this particular PCA decomposition turned out to be, we think it would be instructive when analyzing activity in a network of neurons, to consider at the outset a decomposition in terms of structurally meaningful components, e.g., with a “general spiking” vector, a “left vs right” vector, a “top vs bottom” vector, an “inner layers vs outer layers” vector, etc. or their equivalent depending on the geometry and topography of the brain region under study.

Apart from PCA, we also tested the newer and more complex dimensionality reduction method CEBRA (Schneider et al., 2023). Using the same data as for PCA, we computed a CEBRA embedding in three dimensions (see Methods 2.9). The embedding showed a clear direction of time and the mean trajectories were different for each pattern (Suppl. Fig. 10). The transitions between trajectories for pairs of stimuli were sometimes clear, but not always. In our case we found no clear advantage of CEBRA over PCA.

### Limitations

Notwithstanding its novelty in enabling the study of inter-areal interactions at the level of biophysical mechanisms and connections between realistic neurons, our model has several limitations.

The first major limitation, and perhaps the most evident, is that even though we have two model cortical areas, these are really different volumes extracted from the same cortical area, namely primary somatosensory cortex, and artificially isolated. This means we have the same components and properties (cell morphology and composition, relative layer thickness, local connectivity rules, etc.) in both areas, which would not be the case if they actually represented different cortical areas (Cadwell et al., 2019), especially at different positions along the cortical hierarchy. The model cortical areas do differ, however, in the laminar termination profiles of their long-range connectivity, the presence or absence of sensory inputs, and their local geometry (which has an impact on the topology of local networks, see §2.6 in Reimann et al., 2022). Our model can thus be thought to represent a simplified system but using complex models. The reason for this simplification is twofold. On the one hand, working with isolated volumes connected only through long-range projections is a methodological abstraction to focus on the cortico-cortical loop. On the other hand, it obeys practical considerations, as we did not have another model at the same level of biophysical detail for a different cortical area (e.g., secondary somatosensory cortex), and the production of such a model would have required a huge effort beyond the scope of the present work.

A second major limitation is the fact that we only have two model cortical areas. While we have shown this to be enough to study a one-step cortico-cortical loop and its impact on responses to sensory stimuli, it does not allow to address further topics related to cortical hierarchy, such as: interactions between multiple non-primary sensory areas (i.e., in the middle of the hierarchy) which may be sequential or not (Rockland, 2023), complexification of stimulus features along the hierarchy (as in canonical sensory processing theories, Hubel and Wiesel, 1965), propagation of putative error signals along the hierarchy (as in predictive coding theories, Rao and Ballard, 1999), etc. Also, we are missing in the model other cortical areas (e.g., prefrontal and temporal areas) that play a role in sensory processing as sources of top-down influences (Gilbert & Li, 2013).

The reason for this limitation is, once more, twofold. On the one hand, it is due to how the model was constructed from the base model and its long-range connectome (see Methods 2.3). Finding parts of the base model suitable for extraction as separate cortical areas requires identifying pairs of strongly reciprocally connected locations that are sufficiently apart between them and from the edges to have full disjoint seven-column neighborhoods (the size of our model cortical areas). The existence of such connectivity hotspots is determined by the topographical mapping between somatosensory areas in the base model, which provides limited opportunities of finding them. Thus, we did not attempt to identify a third candidate volume for extraction as a cortical area, and even if we had succeeded in doing so, the other aspect of the limitation would have come into play, which is computational cost.

The running time of our simulations scaled linearly (under the observed sparse firing conditions) with the number of neurons and the duration of the simulation. Adding a third similarly-sized region to the model would have increased the running time of every simulation by at least 50% using the same computational resources, which was beyond practical considering the large number of simulations involved in the present work. For similar reasons, we did not include functional synaptic plasticity in the model.

### Implications for cortical processing

Even if we have naive model that does not implement any particular hypothesis about cortical function, we would like to briefly discuss our results in the context of the main theories of cortical processing. To this end, we refer here to area X as a primary sensory area, and area Y as a higher-order area.

In our model, we have observed the activation of a higher-order area driven by a primary sensory area through feed-forward pathways. This forward step in the cortical hierarchy should involve, according to the canonical theory of cortical processing (Hubel & Wiesel, 1965), a representation of more complex aspects of the stimulus in the higher-order area. The only stimulus-related feature that we observed in the activity of the higher-order area was the specificity of the first active assembly to pairs of stimulus patterns, with a clear topographically defined response to “top” patterns (P_1_ and P_2_) vs. “bottom” patterns (P_3_ and P_4_), as seen in the flat view of the central column (Fig. 3A). While we would not refer to this feature of the stimuli as complex, it represents nevertheless a kind of abstraction (from 4 to 2 alternatives), and it can be readily explained by the underlying connectivity.

We have also observed the activation of a primary sensory area driven by a higher-order area through feedback pathways, both as part of the response sequence from a stimulus presented earlier to the lower-order area, and through direct optogenetic stimulation (Fig. 2B4). In contrast to well-known top-down influences of attentional gain control or increased neuronal tuning (Gilbert & Li, 2013), we observed an approximate linear superposition between the responses to the bottom-up stimulus and to the top-down signal, with practically no change in the response of the stimulus-recipient cells (Fig. 4F left). Furthermore, when presenting competing stimuli we found once more a linear combination of responses (Fig. 5A2), also implying that the top-down signal did not serve as an attention mechanism.

These results could be explained by the aforementioned lack of other cortical areas that would convey specific information associated with task-dependent or behaviorally relevant top-down influences (Gilbert & Li, 2013), or perhaps they are a consequence of the absence of synaptic plasticity in the model, which would underlie perceptual learning and contextual tuning (M. Tsodyks & Gilbert, 2004). The lack of synaptic plasticity would also impede the optimization of prediction errors (Mikulasch et al., 2023), and the observed driving effect of feedback projections is incompatible with the purported suppressive role of top-down predictions (Y. Huang & Rao, 2011). We thus cannot relate our findings to theories of predictive coding. Nevertheless, our model provides a baseline for the integration of stimuli along cortical hierarchies at the smallest level and before being optimized by plasticity.

### Future directions

As has been made evident, one main avenue of future work involves considering the impact of synaptic plasticity. To this end, the model could be readily equipped with the calcium-dependent functional plasticity model (Chindemi et al., 2022) that underlies a plastic version of the base model (Ecker, Santander, et al., 2024). This latter work has demonstrated higher stimulus-specificity of responses after a period of learning, and we would expect to see similar effects in the present model, potentially in both areas. Additionally, it would become possible to learn associations between bottom-up and top-down signals that could lead to behavior beyond the observed linear superposition. It may be the case, however, that functional plasticity is not enough to induce significant changes to the connectome and a model of structural plasticity would have to be incorporated (Iatropoulos et al., 2024). Direct manipulation of the connectome (Pokorny et al., 2024) to make the network less naive could also be an alternative.

Another aspect that could be more or less readily added to the model are neuromodulatory inputs. Previous work on a version of the base model (Colangelo et al., 2022) introduced a model of cholinergic (ACh), dopaminergic (DA) and serotonergic (5-HT) fibers with specific laminar termination profiles, that could be integrated in the present model. This would enable studying the role of these different neuromodulators, variously implicated in attention, arousal and network (de-)synchronization, on sensory processing. Additionally, we could also take into consideration innervation from higher-order thalamus (POm) to L1 and upper L5, that is already built in the model but we did not make use of here.

From an analysis standpoint, we have focused here mainly on the activity of excitatory neurons. It would be worthwhile to look further into the activity of the different populations of inhibitory neurons present in the model, including basket and chandelier cells (PV+), Martinotti cells (SST+) and various types of L1 interneurons (5HT3aR+). Inhibition has been implicated in various aspects of sensory processing, including orientation selectivity (Shapley et al., 2003), negative regulation (feedback inhibition) of primary sensory areas (Hishida et al., 2019) or local gating of information in L1 (S. Huang et al., 2024), among others. Furthermore, we only considered here spiking activity, but the model offers also the possibility of analyzing sub-threshold membrane potential variations across individual neuronal morphologies, as well as the calculation of extracellular potentials (Tharayil et al., 2024).

Looking forward, the most natural elaboration of the present model would be to build a biophysically-detailed model with more cortical areas, where each area would have components and properties particular to that area (cell composition, internal connectivity, etc.). Given the large amount of data required, it would probably be best to base such a model on mouse data, which is readily available nowadays (e.g., Economo et al., 2016; Wang et al., 2020). Beyond the connectivity data employed here (Harris et al., 2019; Knox et al., 2018), increasingly larger electron microscopy datasets (Consortium et al., 2023) could provide increased specificity of long-range connectivity.

On a final note, it has been said that “understanding how different areas of the cortex interact and how the cortex communicates with the rest of the brain is key to deciphering how the cortex performs its functions” (Schuman et al., 2021). We hope to have contributed our grain of sand to this colossal endeavor.

## Data and Code Availability

The model has been released in SONATA format and is freely available on a Zenodo repository (https://doi.org/10.5281/zenodo.14026921) under the CC-BY-NC 4.0 license.

The software used to build the long-range projections in the base model is freely available on a GitHub repository (https://github.com/BlueBrain/white-matter-projections) under the Apache-2.0 license.

## Author Contributions

**Sirio Bolaños-Puchet:** Conceptualization, Methodology, Software, Validation, Formal analysis, Investigation, Data Curation, Writing (original draft, review and editing), and Visualization. **Michael W. Reimann:** Conceptualization, Writing (review and editing), Supervision.

## Supplementary material

**Supplementary Figure 1:**
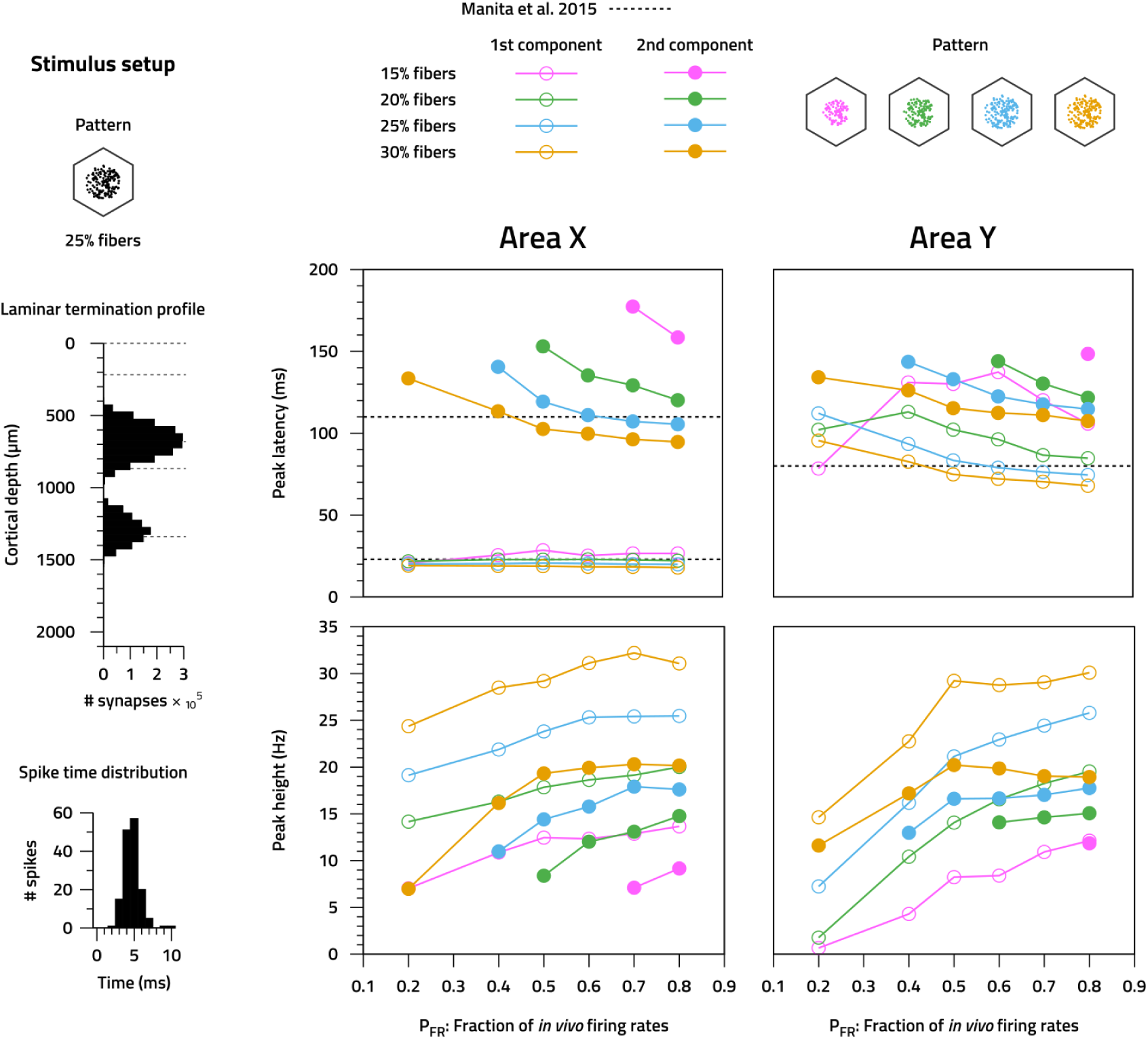
Calibration of stimulus and background activity parameters. Left. Stimulus structure, showing activated fiber pattern (top), stereotypical depth-wise synapse profile (middle), and stereotypical spike time distribution (bottom). Right. Peak latencies (top row) and heights (bottom row) of responses of L5 excitatory neurons in each area (left: area X, right: area Y) to stimuli of various strengths (% of fibers activated) and under various levels of background activity (*P_F R_* parameter). We observe decreasing peak latencies for the second component in area X with increasing stimulus strength and background activity levels, whereas peak heights increase for both components. In area Y a similar behavior is observed.

**Supplementary Figure 2:**
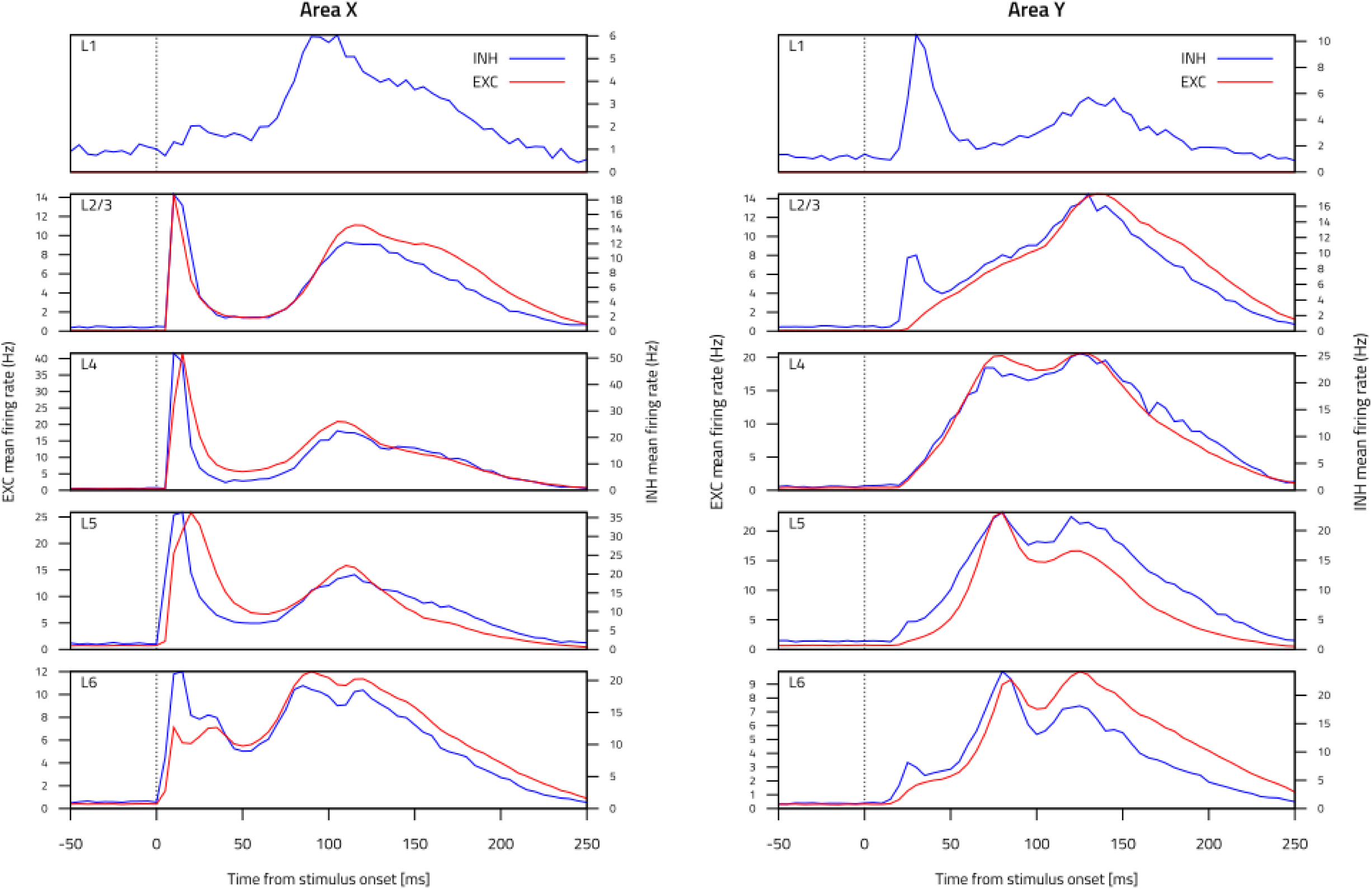
Mean PSTHs across all layer-wise populations. Left. Excitatory and inhibitory populations in each layer of area X: L1, L2/3, L4, L5, L6, from top to bottom. Y axis scaling is specific to each layer, and is different between excitatory (left tics) and inhibitory (right tics) populations. Right. Same, but in area Y.

**Supplementary Figure 3:**
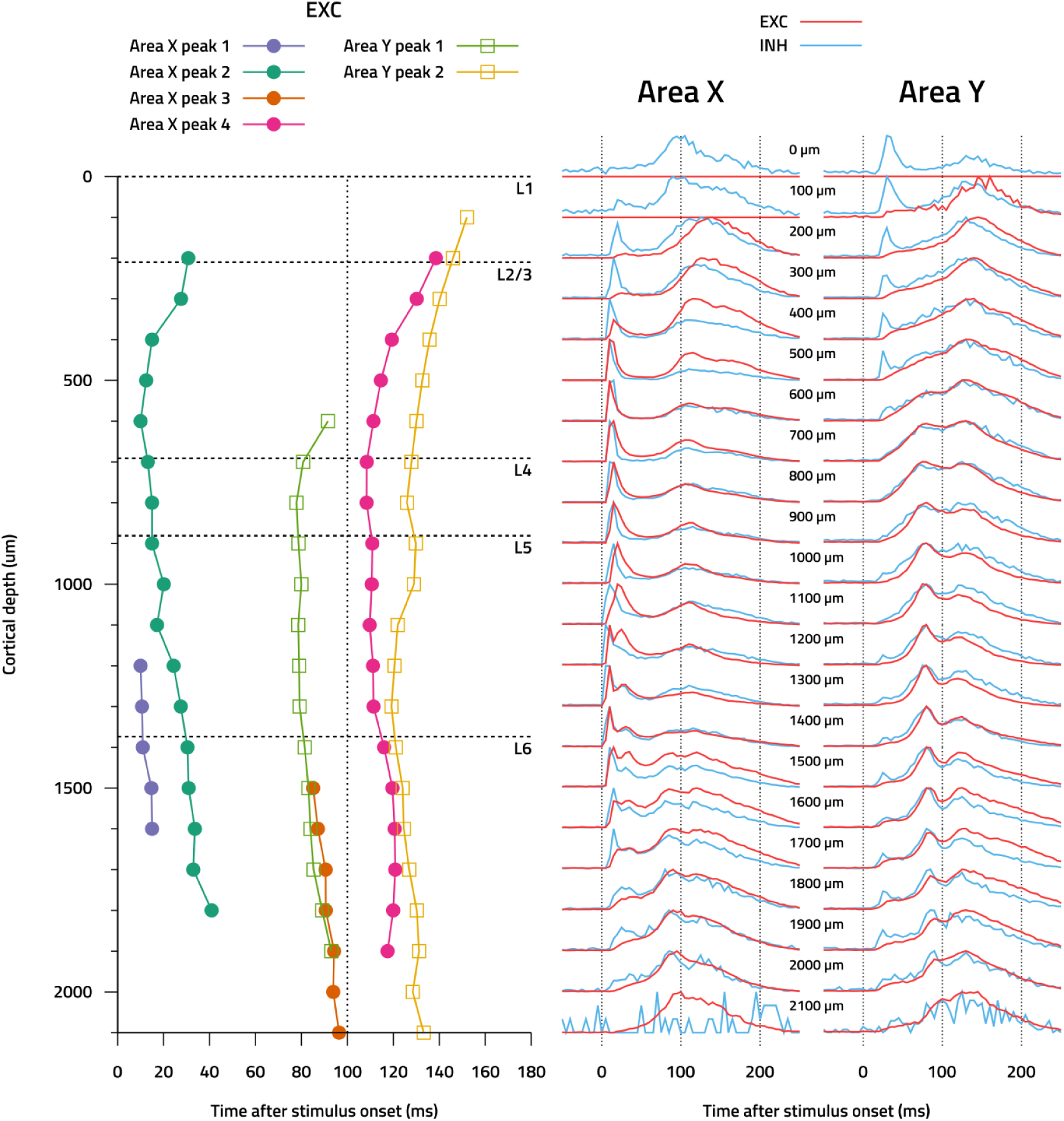
Depth-wise propagation of evoked activity. Left. Peak latencies as a function of depth for all well-defined peaks in the mean PSTHs of excitatory responses in each area. We can observe how activity propagates up and/or down from the point of arrival. Right. Mean PSTHs of excitatory and inhibitory populations in each area (left: area X, right: area Y) as a function of depth (100 *µ*m bins). Subplots have independent scalings of the Y axis to the range of values plotted (also for EXC / INH).

**Supplementary Figure 4:**
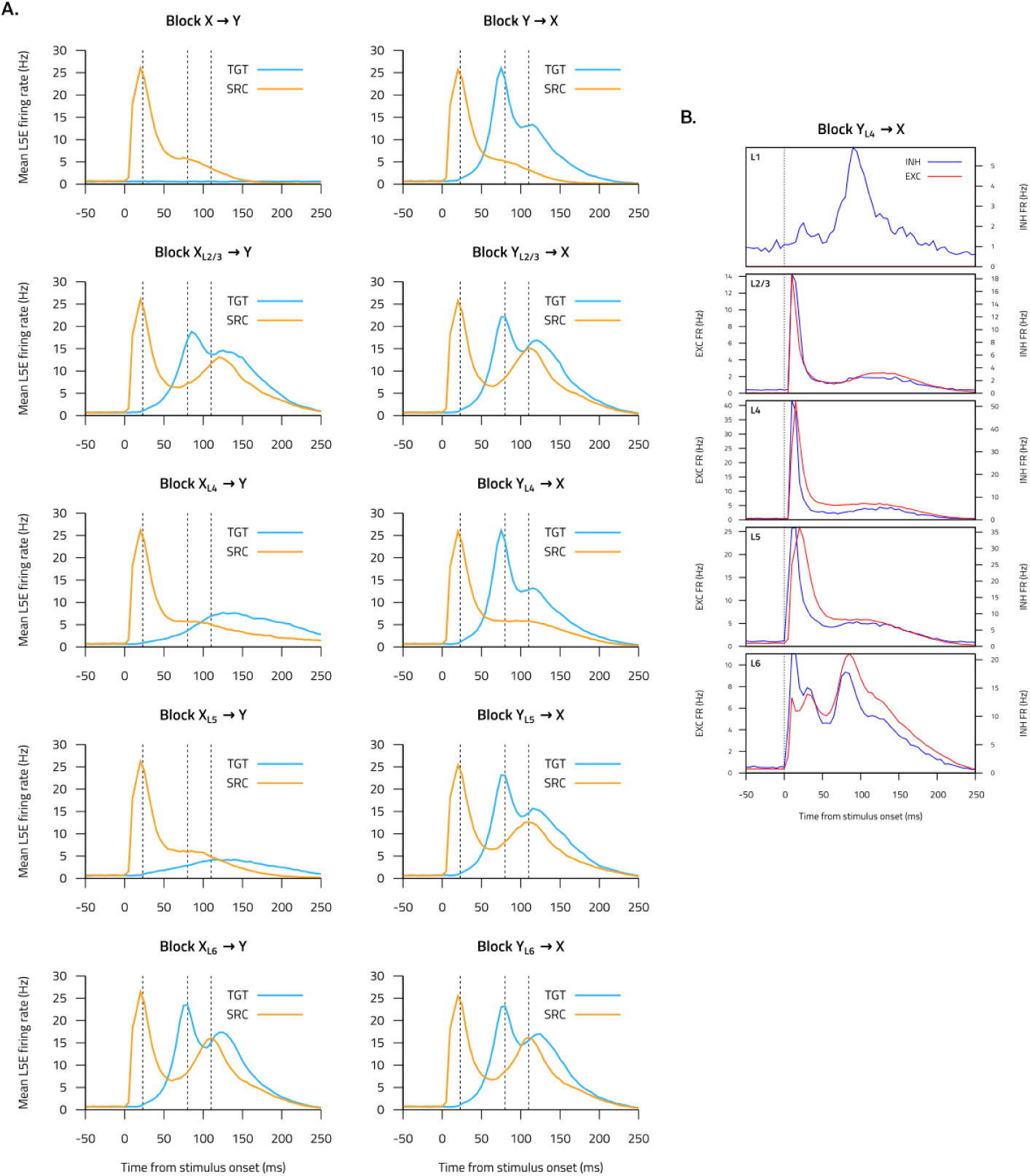
Impact of blocking each layer-wise pathway. **A.** Mean PSTH (N = 30) of layer 5 excitatory neuron responses in area X following application of full or layer-wise blocks in the feed-forward (left column) or feedback (right column) directions. **B.** Mean PSTH (N = 30) of responses in area X across all layers on application of a feedback pathway block from L4 of area Y.

**Supplementary Figure 5:**
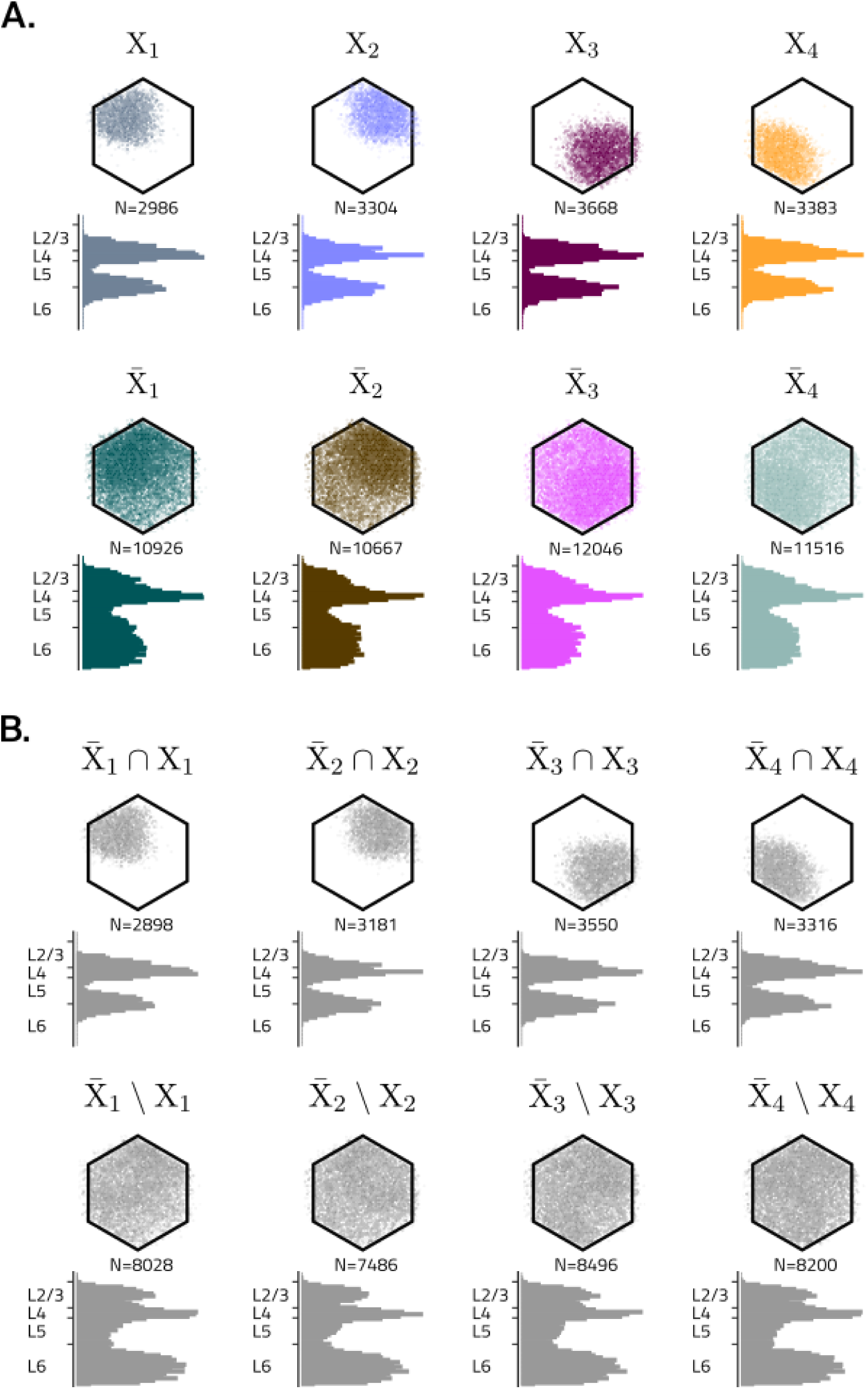
Pattern-specific assemblies in area X and assemblies in area Y. **A.** Locations and depth-wise histograms of neurons in pattern-specific assemblies corresponding to first stimulus presentation (X*_i_*), and second stimulus presentation (X̅ *i*). **B.** Assembly activation sequences in area Y after presentation of single (left column) and paired (right column) stimuli. Arrows indicate the names of the assemblies in the sequences. **C.** Connectivity matrices between assemblies in area Y and pattern-specific assemblies (full and subsets) in area X.

**Supplementary Figure 6:**
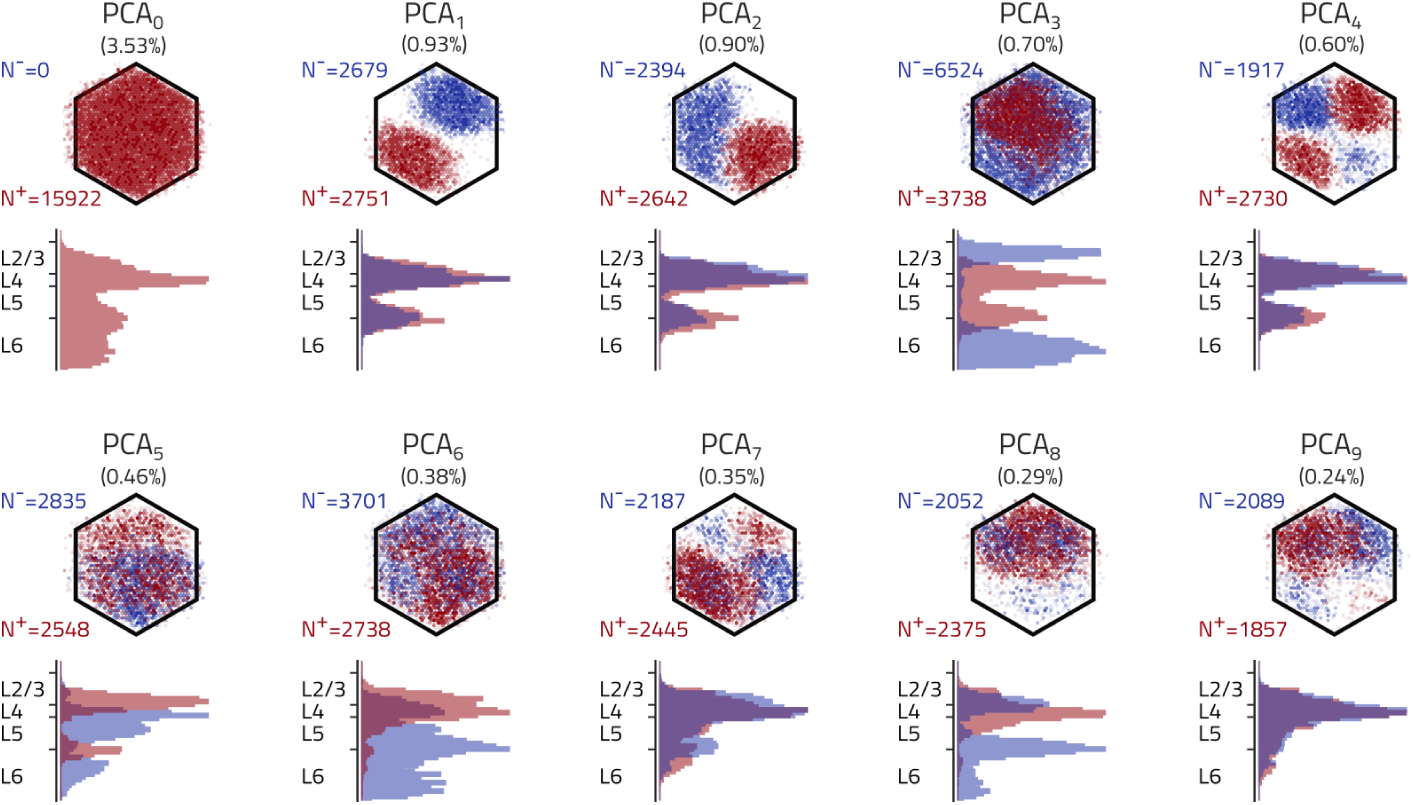
Spatial structure of PCA vectors. Neuron numbers, locations and depth-wise histograms for the first ten PCA vectors. Red/blue are neurons for which the PCA vector has positive/negative components.

**Supplementary Figure 7:**
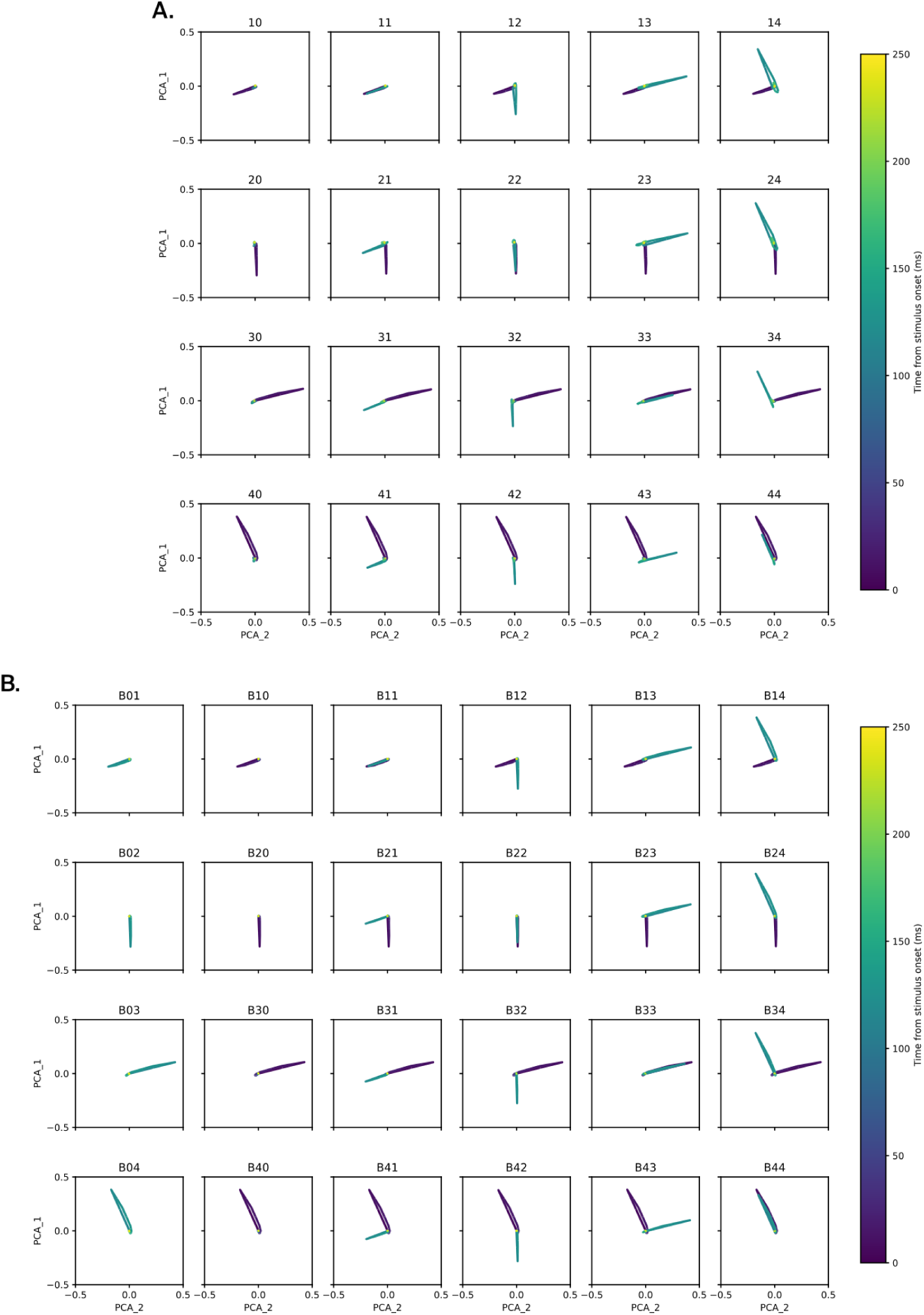
PCA_21_ trajectories across all stimulus pairs. **A.** Network trajectories in PCA_2_-PCA_1_ space for individual patterns and stimulus pairs. Colorbar indicates time along trajectories. **B.** Same, but in the blocked condition. First column is for second stimulus only.

**Supplementary Figure 8:**
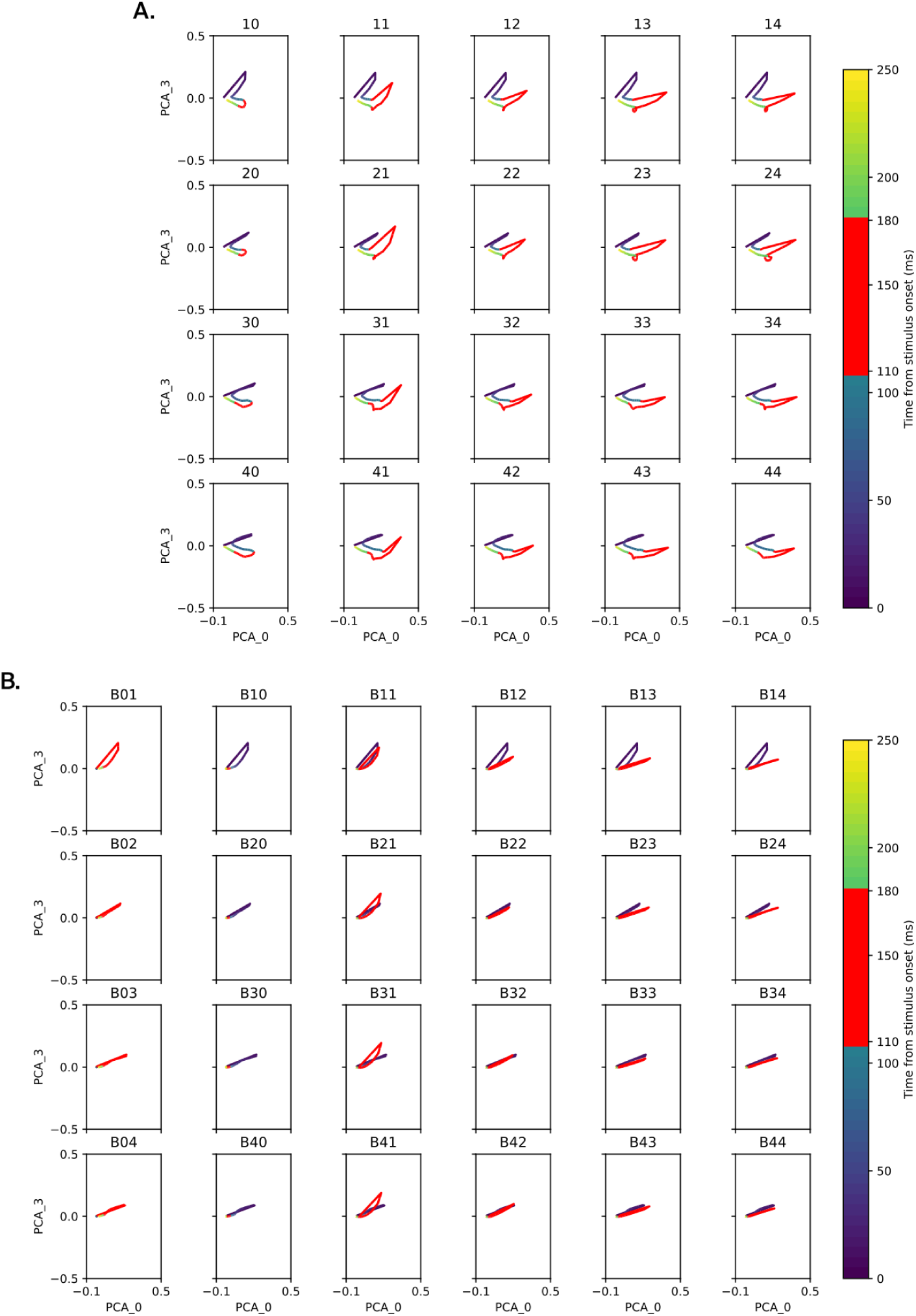
PCA_03_ trajectories across all stimulus pairs. **A.** Network trajectories in PCA_0_-PCA_3_ space for individual patterns and stimulus pairs. Colorbar indicates time along trajectories, with highlight (red) for the time window of feedback interaction (110 to 180 ms after stimulus onset). **B.** Same, but in the blocked condition. First column is for second stimulus only.

**Supplementary Figure 9:**
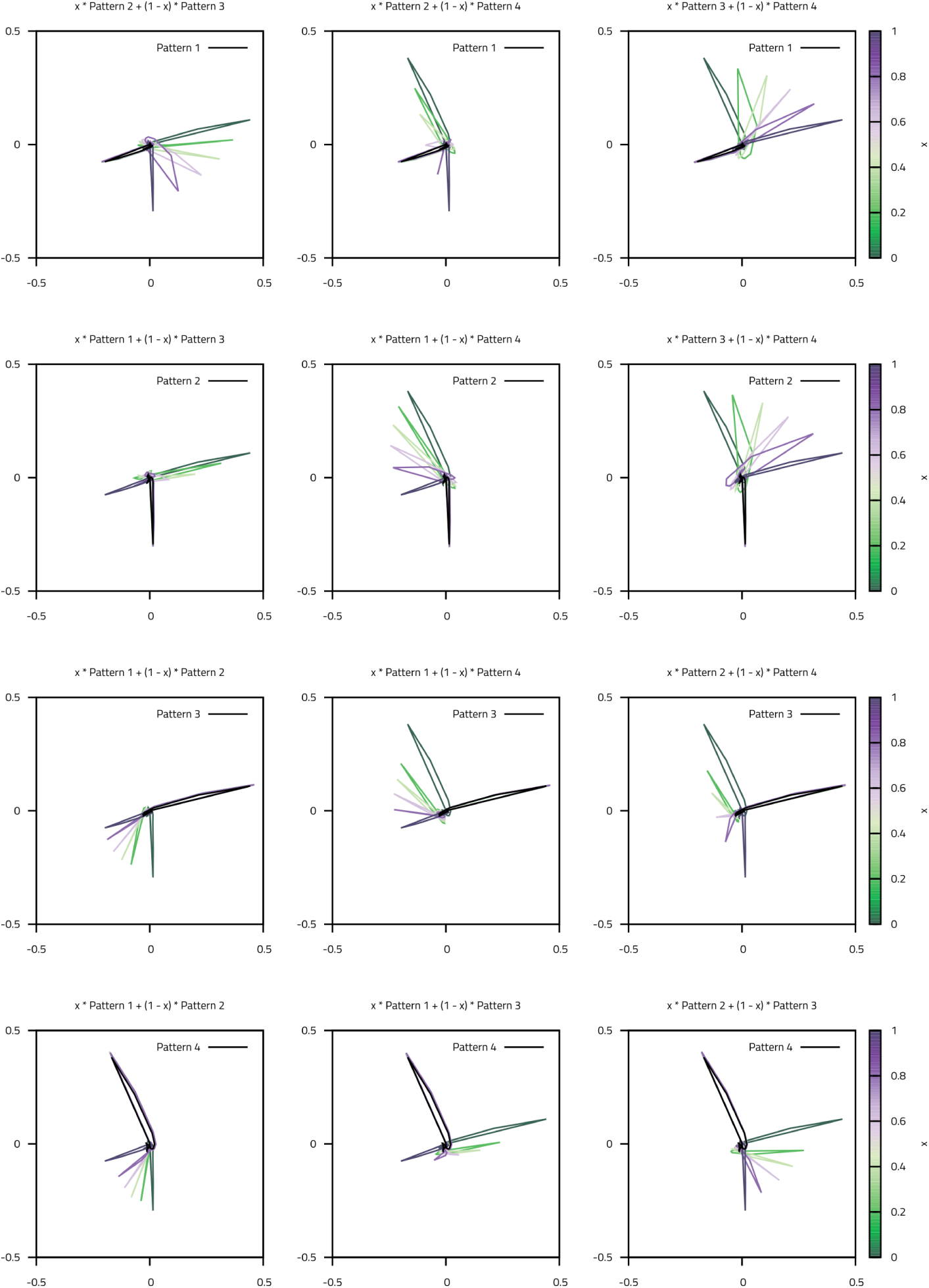
PCA_21_ trajectories across stimulus pairs with interpolated second pattern. Network trajectories in PCA_2_-PCA_1_ space for stimulus pairs where the second pattern is interpolated between two of the primary patterns, i.e. *P_ij_* = *xP_i_* + (1 *− x*)*P_j_*. Colorbar indicates degree of interpolation (*x*).

**Supplementary Figure 10:**
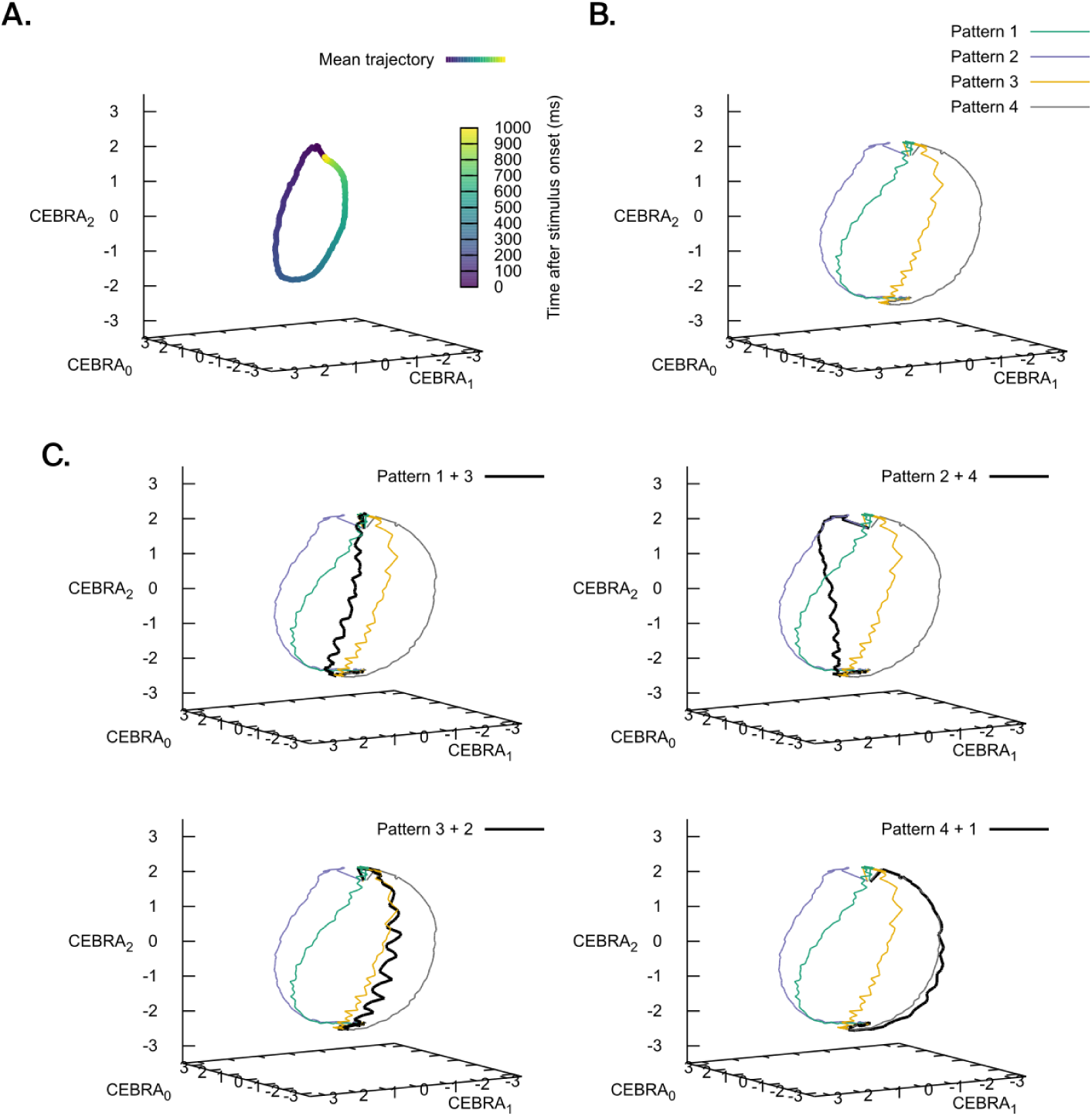
Dimensionality reduction with CEBRA. **A.** Mean network trajectory overall, exhibiting passage of time. **B.** Mean network trajectory of each pattern in the time window from t=0 ms to 400 ms after stimulus onset. **C.** Same as B, with additionally mean network trajectories of example stimulus pairs. We observe the trajectories in the first row starting in the first pattern and moving towards the second of the pair. Trajectories in the second row do not exhibit this behavior, pointing that the embedding better captures the transition from P_12_ to P_34_ than the other way around.

**Supplementary Figure 11:**
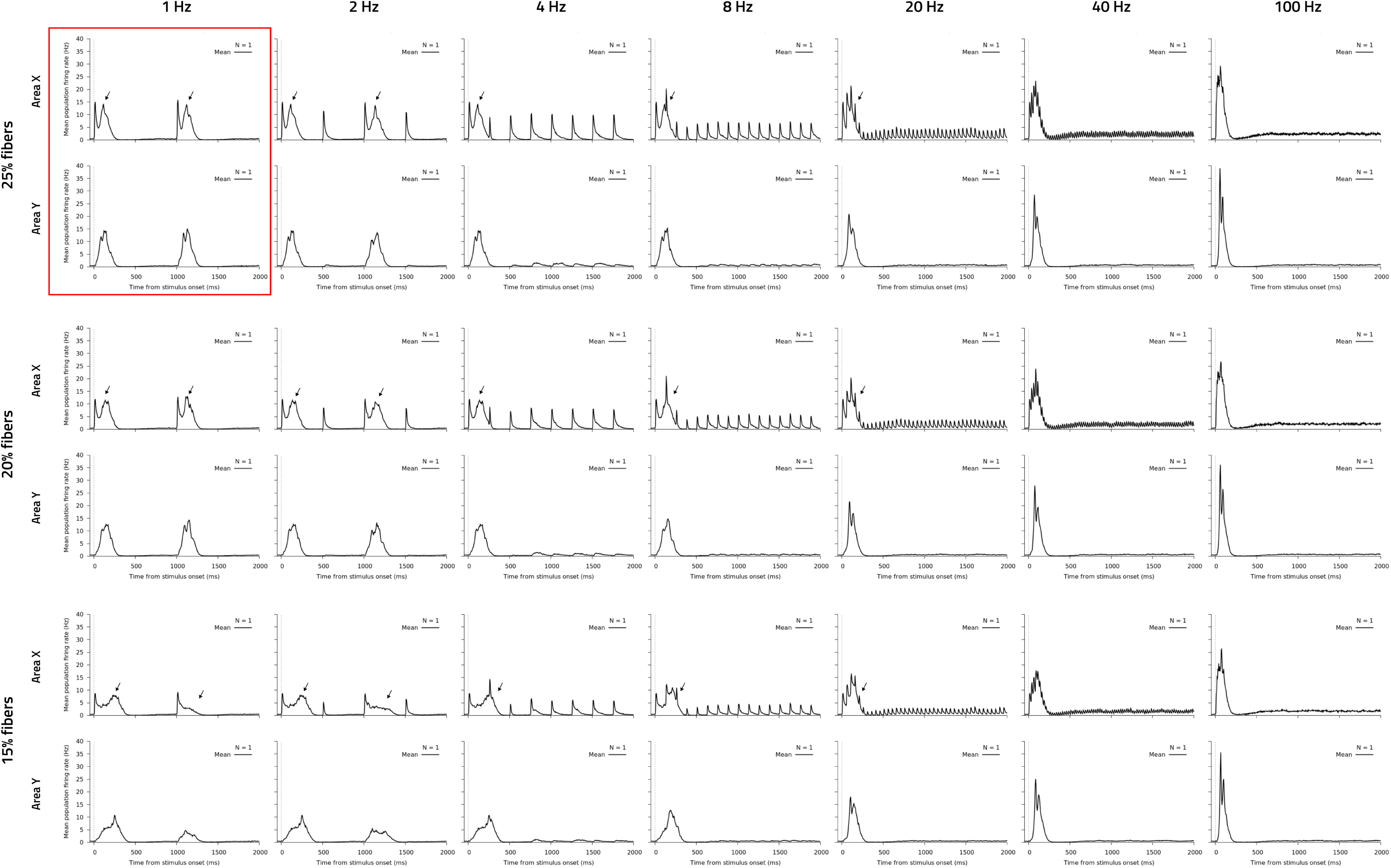
Frequency response of the cortico-cortical loop. Mean population PSTH of responses in both areas (X top, Y bottom) to single stimuli at a given frequency and strength. Columns have increasing frequency to the right (1 Hz, 2 Hz, 4 Hz, 8 Hz, 20 Hz, 40 Hz, 100 Hz). Rows have decreasing stimulus strength values downwards (25% fibers, 20% fibers, 15% fibers). Default case is highlighted (red). Arrows (black) point to clearly visible FD components.

